# Tissue-specific plasticity of DNA methylation across intertidal microhabitats in juvenile mussels *(Mytilus californianus)*

**DOI:** 10.64898/2025.12.12.693252

**Authors:** Qiting Cai, Samuel N. Bogan, Richelle L. Tanner, Joanna L. Kelley, W. Wesley Dowd

## Abstract

Epigenetic modifications to DNA are proposed to underpin plastic responses to environmental change, and the manner in which DNA methylation contributes to plasticity likely differs among tissues. However, few studies have investigated tissue-specific DNA methylation responses to ecologically relevant environmental stressors in natural settings. Here, we used reduced representation bisulfite sequencing on foot and gill to examine the influence of *in situ* microhabitats on DNA methylation in juvenile California mussels (*Mytilus californianus*), a foundation species with widespread dispersal and little evidence of genetic population structure. We examined mussels from a one-month reciprocal transplant experiment between a cool, wave-exposed and a warm, wave-protected microhabitat. These manipulations, which were previously shown to alter juvenile mussels’ heat tolerance, led to significant and highly tissue-specific changes in CpG methylation, including within a number of genes with roles in stress response pathways. Differentially methylated genes were involved in processes including heat shock response, proteolysis, DNA repair, and temperature sensing. In gill, differentially methylated CpGs were more likely to occur in introns relative to other inter- and intragenic features. This study expands on previous research that examined environmentally driven shifts in DNA methylation by documenting plastic and tissue-specific changes in DNA methylation between microhabitats in a natural setting.

## Introduction

The rocky intertidal zone is an exceptionally stressful environment, characterized by extreme wave forces and a dynamic set of other abiotic stressors (Dowd & Denny, In press; Harley & Helmuth, 2003). During tidal cycles, intertidal organisms experience drastic shifts in habitat conditions (Helmuth et al., 2002; Helmuth & Hofmann, 2001), leading to physiological challenges such as temperature extremes, desiccation, and hypoxia (Collins et al., 2023; Helmuth et al., 2006). Climate change will likely exacerbate the frequency and intensity of heat waves and other stressors experienced by many organisms, including those in the intertidal (Sagarin et al., 1999; Stillman et al., 2025). In some instances, these changes may threaten species’ persistence, such as for organisms that currently live close to their thermal tolerance limits (Stillman & Somero, 2000). There is a clear need to better understand the mechanisms used by organisms to respond to present-day challenges as well as their potential to rapidly adjust to future, novel environmental conditions (Hofmann & Todgham, 2010; Somero, 2010; Zhang et al., 2021).

The ability to respond physiologically to environmental fluctuations is particularly critical for the persistence of sessile species, which cannot seek refugia during extreme events. In the rocky intertidal zone, sessile invertebrates play important ecological roles, and many species demonstrate phenotypic plasticity – the ability of a genotype to produce alternative forms of phenotype in a given environment (Padilla & Savedo, 2013; West-Eberhard, 1989) – for physiological traits over various temporal scales when exposed to environmental stressors (reviewed in Leeuwis & Gamperl, 2022). In addition to the temporal variation associated with the tidal cycle, environmental stressors in the intertidal zone can also vary vastly across space.

Exposure to wave action modifies the intensity of intertidal stressors and is one important factor affecting the spatial distribution and abundance of intertidal species (Harger, 1970; McQuaid & Branch, 1985). For example, organisms inhabiting sheltered, wave-protected areas often experience higher thermal stress at low tide than those in wave-exposed habitats (Denny et al., 2011; Hofmann & Somero, 1995). Furthermore, parameters like body temperature can vary at much smaller scales, even among adjacent members of a population (Denny et al., 2011; Helmuth & Hofmann, 2001). This spatial complexity raises the possibility that plastic responses during development (i.e., soon after settlement from the plankton for many intertidal organisms) could help an individual to better match its physiological capabilities with the demands of its adult microhabitat.

Epigenetic changes may mediate some of these plastic responses to the intertidal environment by altering the expression of genes that underlie phenotypic variation (Duncan et al., 2014). Here, we define epigenetics as processes that alter chromatin structure and/or chemically modify nucleotides, thereby influencing gene regulation and ultimately cellular phenotypes (Deans & Maggert, 2015). DNA methylation is one of the most widely studied epigenetic mechanisms and involves the transfer of a methyl group to the fifth carbon of a cytosine residue, specifically at sites called CpG dinucleotides, where cytosine is followed by guanine (Moore et al., 2013). Plastic changes in DNA methylation can regulate gene expression by affecting transcript abundance, alternative splicing, and overall transcriptional homeostasis (Bogan, Strader, et al., 2023; Keller et al., 2016; Li et al., 2018). In some systems, shifts in DNA methylation can modulate the expression of genes in stress response pathways, ultimately contributing to environmental acclimation at the whole-organism level (Bogan & Yi, 2024; Eirin-Lopez & Putnam, 2019). Importantly, the potential gene regulatory roles of DNA methylation likely differ between invertebrates and vertebrates. Vertebrate genomes are known to be extensively methylated, including at promoters whose methylation can silence expression (Keller et al., 2016). In contrast, invertebrate genomes exhibit low levels of methylation, including at promoters, but they are punctuated by regions of increased methylation in gene bodies (Keller et al., 2016; Suzuki & Bird, 2008; Tweedie et al., 1997).

Abiotic environmental stress can induce changes in the set of methylated cytosines of a wide range of organisms, including marine invertebrates inhabiting intertidal and pelagic zones. For example, differential CpG methylation occurs in marine invertebrates upon exposure to ocean acidification (Bogan et al., 2020; Downey-Wall et al., 2020; Liew et al., 2018; Lim et al., 2021; Venkataraman et al., 2022), thermal stress (Guerrero & Bay, 2024; Liu et al., 2023; Roberto et al., 2021; Wang et al., 2021), and varying salinity levels (Johnson et al., 2022; Zhang et al., 2017). Differences in DNA methylation may also contribute to phenotypic diversity across space in natural settings. In one widely-dispersing species, epigenetic divergence was found to be greater than genetic divergence among eastern oyster *Crassostrea virginica* populations from the Gulf of Mexico (K. M. Johnson & Kelly, 2020). At smaller spatial scales, divergent DNA methylation signatures were observed between intertidal and subtidal zones in both Antarctic limpets (*Nacella concinna*; Clark et al., 2018) and Pacific oysters (*Crassostrea gigas*; Wang et al., 2021, 2023). Importantly, studies of epigenomic variation between populations or across intertidal microhabitats have not determined how much of this variation is attributable to plastic changes in methylation, as opposed to prior microhabitat sorting or other processes. Reciprocal transplant experiments combined with epigenetic analyses are a potentially powerful approach to detect differential methylation (DM) that is linked to plastic responses to the environment (Chapelle & Silvestre, 2022; Clark et al., 2018). However, the full potential of transplant experiments to inform ecological epigenetics has not been realized (Bossdorf et al., 2008; Johnson et al., 2022; Sork, 2018).

Here, we pair reciprocal transplant experiments with DNA methylation analysis to examine environmentally driven plasticity among juveniles of the California mussel (*Mytilus californianus*), a competitive dominant intertidal bivalve found along the western coast of North America (Paine, 1974; Seed & Suchanek, 1992). Mussel beds play an important ecological role by supporting plant, animal, and microbial diversity in the intertidal zone (Pfister et al., 2014; Seed & Suchanek, 1992). *M. californianus* has a long planktonic larval duration, long dispersal distances, and high levels of gene flow among populations, each contributing to low population genetic structure along its range (Addison et al., 2008; Levinton & Suchanek, 1978; Lopez-Duarte et al., 2012). Plastic responses to environmental change have been recorded in *M. californianus* from molecular to whole-organism levels and across life history stages (Buckley et al., 2001; Gleason et al., 2023; Jimenez et al., 2015; Williams & Somero, 1996). For instance, at the molecular level, individuals’ thermal histories influenced the induction temperature for heat-shock protein production (Buckley et al., 2001). A reciprocal transplant study found that newly settled juvenile mussels exhibited increased upper thermal tolerance after transplantation from cool (wave-exposed) to warm (wave-protected) microhabitats for one month in summer (Gleason et al., 2018). It was hypothesized that epigenetic modifications, such as DNA methylation, may have underpinned the observed pattern of developmental plasticity in heat tolerance (Gleason et al., 2018).

Using samples collected from juvenile mussels from Gleason et al. (2018), we first evaluate the prediction that transplantation to a warmer, wave-protected field site induces a distinct pattern of DNA methylation. We further explore how the difference in wave exposure between microhabitats alters DNA methylation of juvenile mussels in a tissue-specific manner, using gills and foot. Studies of plastic responses to the environment tend to focus on the performance of whole organisms or on molecular responses in a single tissue (reviewed in Husby, 2020), but the latter approach could bias interpretation and/or miss critical responses in tissues that were not sampled. For example, tissue-specific gene expression and lipidomic responses to laboratory-based experimental warming, salinity stress, and pollutant exposure were observed in the gills, digestive gland, and mantle in some *Mytilus* mussels (Guinle et al., 2025; Wu et al., 2022). These issues are particularly relevant for studies of DNA methylation, which in many organisms including molluscs exhibits tissue-specific patterns associated with tissues’ distinct biological functions (Geyer et al., 2017; Haidar et al., 2024; Li et al., 2015). In bivalve molluscs, the gills use cilia to generate feeding currents and serve as the primary site for gas exchange (Gosling, 2015). Given their large surface area and these roles as an interface between organisms and the environment, gills are one of the most metabolically active tissues and are sensitive to environmental stressors, including temperature (Châtel et al., 2011; Dahlhoff et al., 2002; Jimenez et al., 2015; Lockwood & Somero, 2011; Tomanek & Zuzow, 2010). Gills can also be prone to oxidative damage under environmental stress (Almeida et al., 2005; Wang et al., 2018). The foot, in contrast, is a muscular tissue important for body positioning. In mussels, the foot also secretes byssal threads, which are essential for attaching to substrates and preventing dislodgement by wave motions (Carrington et al., 2008). Thermal stress can lead to foot structural abnormalities and reduced adhesion performance (Li et al., 2020; Park et al., 2015). Considering the disparate functions of gills and foot, we predict that these tissues might also exhibit distinct epigenetic responses to changes in the abiotic environment. However, most of the existing studies of DNA methylation responses to environmental stressors in invertebrates have only focused on a single tissue, such as gills (e.g. Johnson et al., 2022; Roberto et al., 2021; Wang et al., 2021; Zhang et al., 2017) or mantle (e.g. Downey-Wall et al., 2020). Furthermore, field-based studies investigating tissue-specific molecular responses to environmental stressors exist across taxa but are limited (e.g., Escobar-Sierra et al., 2024; Pereira et al., 2025; Rossi et al., 2016; Watson et al., 2017). The role of tissue-specific DNA methylation adjustments in responding to natural environmental changes remains largely unexplored (Haidar et al., 2024); to our knowledge, no *in situ,* multiple-tissue epigenomic studies have been reported.

In summary, we use reduced representation bisulfite sequencing (RRBS) to identify methylated CpG loci and to compare levels of DNA methylation between juvenile mussel tissue types and among reciprocal transplant treatment groups. We predicted that the one-month reciprocal transplant would lead to different DNA methylation signatures between the two tissues in genes associated with stress responses, and that differential methylation after transplantation would be more pronounced in gills due to their high metabolic demand and environmental sensitivity (Dahlhoff et al., 2002; Dowd et al., 2013; Gosling, 2015). We also predicted that DNA methylation patterns in the two tissue types would be shaped by both the transplant treatments (i.e., acclimatization to the most recently experienced conditions) and the original sites at which the mussels first settled out of the plankton (i.e., habitat sorting and/or post-settlement selection). The results will help identify potential non-genetic mechanisms by which juvenile mussels regulate their physiology under the demands of unpredictable post-settlement environments and how they might respond to environmental stressors in the face of future climate changes. More generally, this study extends our knowledge of the drivers of epigenomic diversity in nature by evaluating environment-associated differences in methylation over small, microhabitat scales and by exploring whether environment-methylation associations during development differ between tissue types. Understanding tissue-specific epigenetic responses to environmental changes *in situ* is important for developing a more comprehensive understanding of how organisms will cope with the challenges presented by climate change.

## Methods

### Reciprocal transplant experiment

All tissue samples analyzed in this study were collected at the conclusion of the reciprocal transplant experiment conducted by Gleason et al. (2018); that experiment is briefly summarized here. Juvenile *M. californianus* were originally collected from either high- or low-shore mussel beds at both wave-exposed (cool) and wave-protected (warm) sites at the Hopkins Marine Station in Pacific Grove, CA, USA (36.62178 °N, 121.90438 °W) in June 2016. Mussels from each of these ‘origin’ sites were transplanted to a low-shore location at either their original wave-exposure condition or the opposite condition during early summer of 2016 (n = 562). iButton temperature data loggers (model DS1921G, Maxim Integrated, San Jose, CA, USA) embedded in silicon-filled adult mussel shells were used to assess thermal differences between the transplant sites. The resulting average daily range of body temperatures was three times greater at the protected site compared to the exposed site (8.1°C vs. 2.7°C). During the transplantation period, individuals were placed inside a mesh bag, enclosed within a stainless-steel antipredator cage (3.3 mm mesh), and attached to an acrylic plate that was bolted to the rock substrate. They were retrieved after one month (July; Gleason et al., 2018). Shell lengths were measured immediately before and after the transplant experiment with digital calipers to determine growth rate and length at time-of-sampling (Gleason et al., 2018). The mussels sampled for the present study were not phenotyped for heat tolerance at the conclusion of the original transplant treatments, because that acute heat stress might have impacted epigenetic patterns. Instead, the tissues were flash frozen in liquid nitrogen.

All statistical analyses were performed in *R* v4.3.2 (R Core Team, 2023). Only the final shell length measurements from July were included in analyses. Because juvenile mussels collected from high-shore or low-shore locations exhibited very similar phenotypic responses in terms of the change in heat tolerance (Gleason et al., 2018), in the present study, we only examined the effect of origin site wave-exposure (exposed or protected) and transplant site wave-exposure (exposed or protected) on methylation patterns of mussels from the two low-shore origin sites.

### Reduced representation bisulfite sequencing (RRBS)

A total of 65 tissue samples from 39 juvenile *M. californianus* were dissected from gill and foot. DNA was extracted from the tissue samples for bisulfite sequencing. Of these, 52 samples were collected from the same 26 individuals. Using the Qiagen Puregene Tissue Core Kit A, genomic DNA was isolated separately from each tissue type for each sample. Following a modified version of the Gu et al. (2011) protocol for generating RRBS libraries, an *MspI* digestion (NEB Catalog #R0106) was performed with a phenol-chloroform-isopropyl (3:1:1) clean up. *MspI*-digested DNA was end-repaired and A-tailed using a Klenow fragment (NEB Catalog #M0212), and once again cleaned with phenol-chloroform-isopropyl (3:1:1). A-tailed (NEB custom order) DNA was ligated with methylation-specific NEBNext adapters, and a 1.5X SPRI bead clean-up was performed (Omega Mag-Bind Total Pure NGS magnetic beads). Then, bisulfite conversion of ligated DNA was performed with the EZ DNA Methylation-Gold Kit (Zymo Research, catalog D5005 and D5006). DNA was amplified while simultaneously attaching NEBNext Multiplex Oligos for Illumina (unique dual index primer pairs, Catalog #E6440) using 16 PCR cycles: 95°C for 2 min to denature; then 16 cycles of 95°C for 30 s (denature), 65°C for 30 s (anneal), 72°C for 45 s (extend); then 72°C for 7 min. A final 1.8X SPRI bead clean-up for the pooled libraries was then performed. Pooled libraries were sequenced on four lanes for each run on an Illumina HiSeqX (150 base pair paired end) at the University of Southern California Genome Core.

### Quality assessment, trimming, mapping, and methylation calling

Sequence quality of raw reads was assessed with FastQC v0.12.1 (Andrews, 2010) and MultiQC v1.25.2 (Ewels et al., 2016). Raw RRBS reads were trimmed using TrimGalore! v0.6.10 (https://github.com/FelixKrueger/TrimGalore) with the --rrbs and paired-end option. To reduce methylation bias due to an increase in methylation calls at the 3’ ends of Read 2, all the paired-end reads were hard trimmed to only retain 65 base pairs from the 5’ end of the reads. Sequence quality was assessed again with FastQC v0.12.1 (Andrews, 2010) and MultiQC v1.25.2 (Ewels et al., 2016) after trimming. For each sample, trimmed reads from each of the four lanes of the same sequencing run were concatenated. The concatenated reads were aligned to the *M. californianus* reference genome xbMytCali1.0.p (GCF_021869535.1; Paggeot et al., 2022) using Bismark v0.24.2 (Krueger & Andrews, 2011) with the paired-end option and a minimum score threshold of ‘L, 0, -0.6’ and a stringency of 1. The reference genome was first bisulfite converted for C->T and G->A using the command bismark_genome_preparation. Methylation calls were extracted using the command bismark_methylation_extractor. Since our hypotheses were not specific to strandedness, both strands of the methylation coverage files produced by bismark_methylation_extractor were merged following recommendations by Venkataramann and colleagues (Bogan, Johns, et al., 2023; Venkataraman, Greiner-Ferris, et al., 2024).

### Annotation of features containing CpGs

The genomic features where CpG sites were located were examined using the *R* packages *genomation* v1.34.0 (Akalin et al., 2015) and *GenomicRanges* v1.54.1 (Lawrence et al., 2013). The annotation .gff file from the reference genome xbMytCali1.0.p (GCF_021869535) was downloaded from NCBI. Introns and Untranslated Region (UTR) annotations were added to the original annotation using the command line tool *agat* v1.2.0 (Dainat, 2020). Promoter regions were defined as 1 kilobase upstream of the first exon of each gene. Intergenic regions were identified as regions that did not fall into exons, introns, UTRs, or promoters. Each CpG site was categorized based on genomic position: intron, exon, promoter, intergenic, and/or UTR. If a CpG site was in multiple genomic features (e.g., intron and exon of different genes), all features were retained in the analysis.

### Analysis of multivariate CpG methylation patterns using principal component analysis (PCA)

To enable exploration of the effects of origin site, transplant site, tissue, and other variables on patterns of methylation, a methylation proportion matrix that included all 65 foot and gill samples was first generated. Methylation proportion was calculated as methylated reads / (methylated reads + unmethylated reads) per CpG site per sample. *mice* v3.16.0 (Van Buuren & Groothuis-Oudshoorn, 2011) was used to impute methylation proportions at CpGs that were missing in a sample.

To assess whether multivariate CpG methylation clustered according to tissue type, origin site, and/or transplant site, a principal component analysis (PCA) was performed on the imputed methylation proportion matrix of all samples using the prcomp function from the *R* package *stats* v4.3.2. To examine the influences of tissue type and final shell length on the PCA loadings in this genome-wide analysis, we then ran a linear mixed-effect model that included sample ID as a random effect using the *lmer* function from *R* package *lme4* v1.1-35.5 (Bates et al., 2015). Lastly, we assessed whether foot and gill samples collected from the same individuals clustered more closely together in the PCA space than tissues from different individuals. To this end, a Mantel matrix correlation test was performed using the *mantel* function from the *R* package *vegan* 2.6-10 (Oksanen et al., 2025). The correlation between a matrix of inter-sample methylation distances versus a binary identity matrix was assessed. Methylation distance was measured as Euclidean distance across PC axes 1 – 19, which captured 51.0% of the total variance of the data. Distances in the binary identity matrix were set to 0 for tissues of the same individual, and 1 for samples from different individuals.

### Genome-wide methylation levels

The full dataset was further filtered for all subsequent analyses. First, to prevent spurious model predictions, significant outlier samples identified by the *arrayQualityMetrics* distance-between-arrays test in *R* were removed (Kauffmann et al., 2009; Supplemental Materials). This removed three significant outlier gill samples (15W-G_S12, 60G-G_S58, 38B-G_S78); no outliers were detected among foot samples. To further maximize the number of CpGs retained in the filtered dataset, the five remaining samples with the largest library sizes from each origin-by-transplant treatment group for each tissue were selected for all downstream analyses (n = 40, 20 each for foot and gill tissue; Table S1). Of these samples, 20 were paired foot and gill tissues collected from the same 10 individuals, and the remaining 10 foot and 10 gill samples came from unique individuals (Table S1).

As an additional quality control check of the RRBS libraries, the percent methylation of reads aligned to CpGs was compared between tissues and treatment groups (see Supplemental Methods). This analysis of methylation in CpG-aligned reads ensured that (i) samples or treatment groups did not differ in RRBS coverage in methylated regions and (ii) samples exhibited comparable levels of bisulfite conversion.

We then tested whether treatment groups and tissues exhibited differences in genome-wide CpG methylation. To achieve this, we calculated the mean proportion of methylation across all CpGs per sample in the filtered dataset, using only CpGs with at least three reads in 14 of the 20 samples per tissue type. A generalized linear mixed model with a beta distribution and logit link were fitted with *R* package *glmmTMB* 1.1.10 (Brooks et al., 2017). This model predicted variation in genome-wide mean CpG methylation as a function of fixed effects for tissue type, origin site, and transplant site. Sample ID was included as a random effect to control for non-independence between tissue samples from the same individual.

### Differential methylation analysis

Differential methylation (DM) analysis was performed with the *R* package *edgeR* v4.0.16 (Robinson et al., 2010) using coverage files generated from Bismark (Krueger & Andrews, 2011). Use of *edgeR* for DM analysis of RRBS data followed the recommendations of Chen et al. (2018). Separate models of DM were fitted to the filtered CpG methylation matrices from the 20 gill and 20 foot samples. The *edgeR glmFit* function was used to fit negative binomial generalized linear models that predicted variation in methylated and unmethylated read counts as a function of fixed effects for origin site, transplant site, and the shell length of the mussels at time of sampling (Robinson et al., 2010). Shell length was included as a covariate because of its correlation with sample loadings to methylation PC1 in the PCA analysis (see below). We also evaluated the performance of a more complex model that was fitted with the same fixed effects and an interaction term between origin and transplant site. However, this model risked overfitting compared to the iteration lacking the transplant x origin interaction; details are provided in the Supplemental Materials. Differentially methylated CpGs (DM CpGs) associated with transplant site, origin site, and shell length were identified after adjusting for multiple hypothesis testing to control the false discovery rate (FDR) using the Benjamini and Hochberg method (Benjamini & Hochberg, 1995) and an FDR alpha value of < 0.05.

The associated log2-fold-change (log2FC) for each DM CpG represents an increase or decrease in methylation in the wave-protected relative to the wave-exposed groups, for both the origin and transplant factors. These log2FC values were further analyzed for directional bias and tissue differences. First, two-sided Wilcoxon signed-rank tests using the *wilcox.test* function in *R* were performed on the log2FC values associated with both the origin and transplant site effects to determine whether methylation changes were significantly skewed toward hyper- or hypomethylation in each tissue. Linear models were then fitted to assess whether the directionality and overall strength of differential methylation were significantly different between treatment groups and tissue types by comparing the mean log2FC of significant DM CpGs (and absolute log2FC) using the *lm* function in *R*. Lastly, associations between all CpG log2FC values (regardless of significance) shared between the two tissues for origin and transplant site effects were evaluated using Pearson’s correlation tests with the *cor.test* function in *R*.

We tested for over- or underrepresentation of DM CpGs within major classes of genomic features (introns, exons, intergenic regions, promoters, and UTRs) relative to genomic background. This was achieved by comparing the numbers of DM CpGs in each genomic feature to the distribution of all available CpG sites across these features genome-wide with two-sided Fisher’s exact tests using the function *fisher.test* in *R*. Fisher’s exact tests were conducted separately for transplant and origin effects in each tissue type. Fisher’s exact tests were also used to compare the distribution of DM CpGs across genomic features between tissues. FDR correction was applied to p-values from all Fisher’s exact tests using the Benjamini and Hochberg method (Benjamini & Hochberg, 1995). This approach identified features that were over- and underrepresented for DM relative to background, but it did not determine whether differences in DM among feature types were significant. Thus, significant differences in the prevalence of DM CpGs among genomic features were also evaluated using generalized linear models (GLMs) with a binomial distribution and logit link using the *glm* function in *R* (Supplemental Materials).

### Gene Ontology (GO) enrichment analysis

Gene Ontology (GO) enrichment analysis was performed separately for each tissue type to investigate the functions of the DM genes using the *R* package *GOseq* v1.54.0 (Young et al., 2010). DM genes were defined as genes that contained at least one significant DM CpG site and at least three CpGs in the coverage filtered dataset within the gene region. A rank-based Gene Ontology Analysis with Adaptive Clustering and Mann-Whitney U test based on ranking of mean gene-wide log2FC of DM (GO-MWU, https://github.com/z0on/GO_MWU) was also conducted. Detailed methods for the GO enrichment analyses can be found in the Supplemental Materials. Because GO annotations are both incomplete and biased toward reporting of functions in humans and other model organisms, we supplemented the GO analysis of differentially methylated genes with manual review of their potential roles in stress response pathways based on previously reported findings in the literature.

## Results

### PCA analysis revealed little effect of origin, transplant, or tissue on genome-wide methylation patterns

Methylated and unmethylated CpGs in the full dataset of 65 mussel samples were identified from the Bismark alignment outputs (Table S2). As expected for an invertebrate, the mean genome-wide CpG methylation was low (<15%; Table S2). A Principal Component Analysis (PCA) of the DNA methylation proportions across all samples and all CpG sites did not separate samples by tissue type, origin site, or transplant treatment (Figure 1A). The top two principal components explained a combined 6.78% of variation in CpG methylation. Shell length exhibited a significant and positive correlation with PC1 (marginal R^2^ = 0.37; p < 0.001; Figure 1B, Table S3a). No significant relationship was detected between shell length and PC2 (Figure S1, Table S3a). The shell lengths of mussels ranged from 7.75mm to 15.02mm (mean ± SD: 10.74 ± 2.17mm; Table S1).

**Figure 1.**
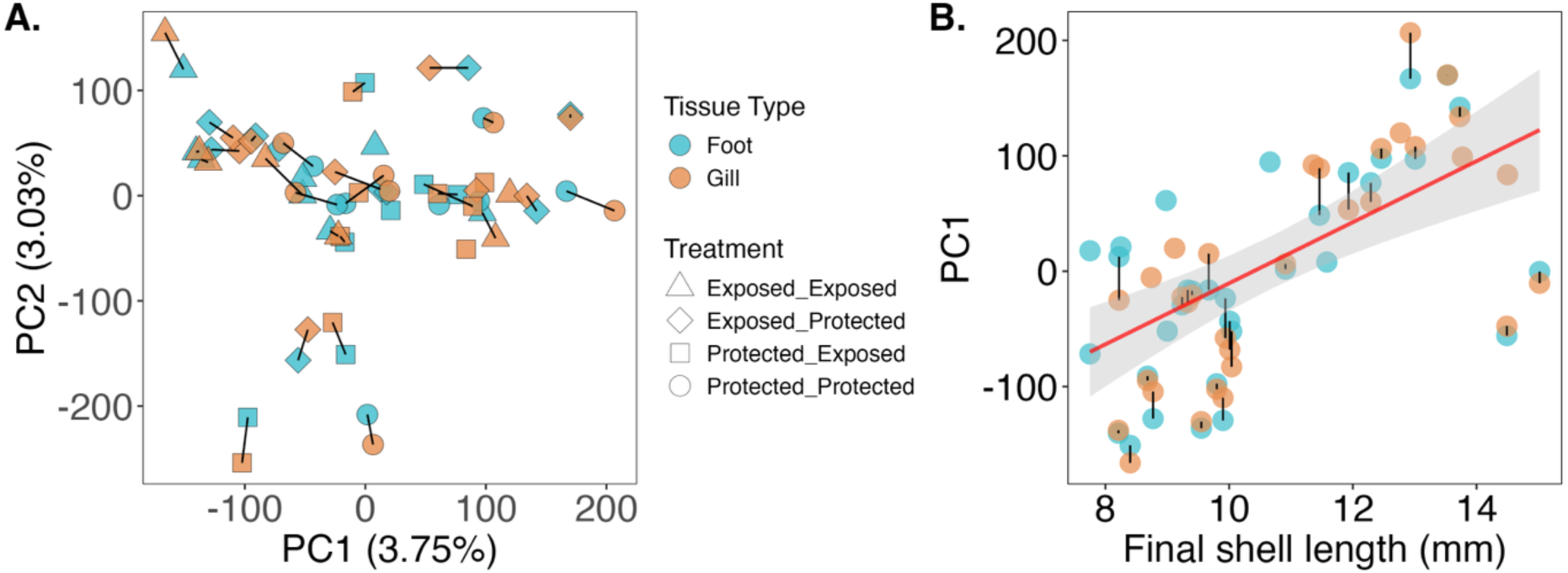
Principal Component Analysis (PCA) on the full matrix of CpG methylation data. Methylation proportion was calculated as methylated reads/(methylated + unmethylated reads). Imputations were created for CpG sites with missing data. Tissues collected from the same individual are connected with a black line; N = 33 for foot and N = 32 for gill. **A**. PCA performed on proportions of methylation across all CpGs did not separate samples by tissue type or treatment. Shapes correspond to origin and transplant treatment. **B**. a positive correlation was found between PC1 and shell length (marginal R^2^ = 0.37; p < 0.001). The red line represents the best-fit line, and the ribbon represents the 95% confidence interval.

Notably, foot and gill tissues collected from the same individual appeared to cluster closely together in PC space (Figure 1A). A Mantel test showed a significant, positive correlation between a matrix of Euclidean methylation distances in PC space (using scores on the first 19 principal component axes) and a binary matrix of mussel identity (Mantel r = 0.33, p = 0.001).

### Genome-wide CpG methylation levels

After removing outliers and retaining only the five samples with the largest library sizes in each treatment-by-tissue combination, we calculated mean genome-wide methylation levels from Bismark outputs (% methylation of reads in CpG context). This metric did not differ between tissue types, origin sites, or transplant sites (p > 0.05 for each; Figure S2, Table S3b). The same pattern was observed when genome-wide methylation was calculated as mean methylation proportion across CpG loci after coverage filtering (p > 0.05 for each; Figure 2, Table S3b).

**Figure 2.**
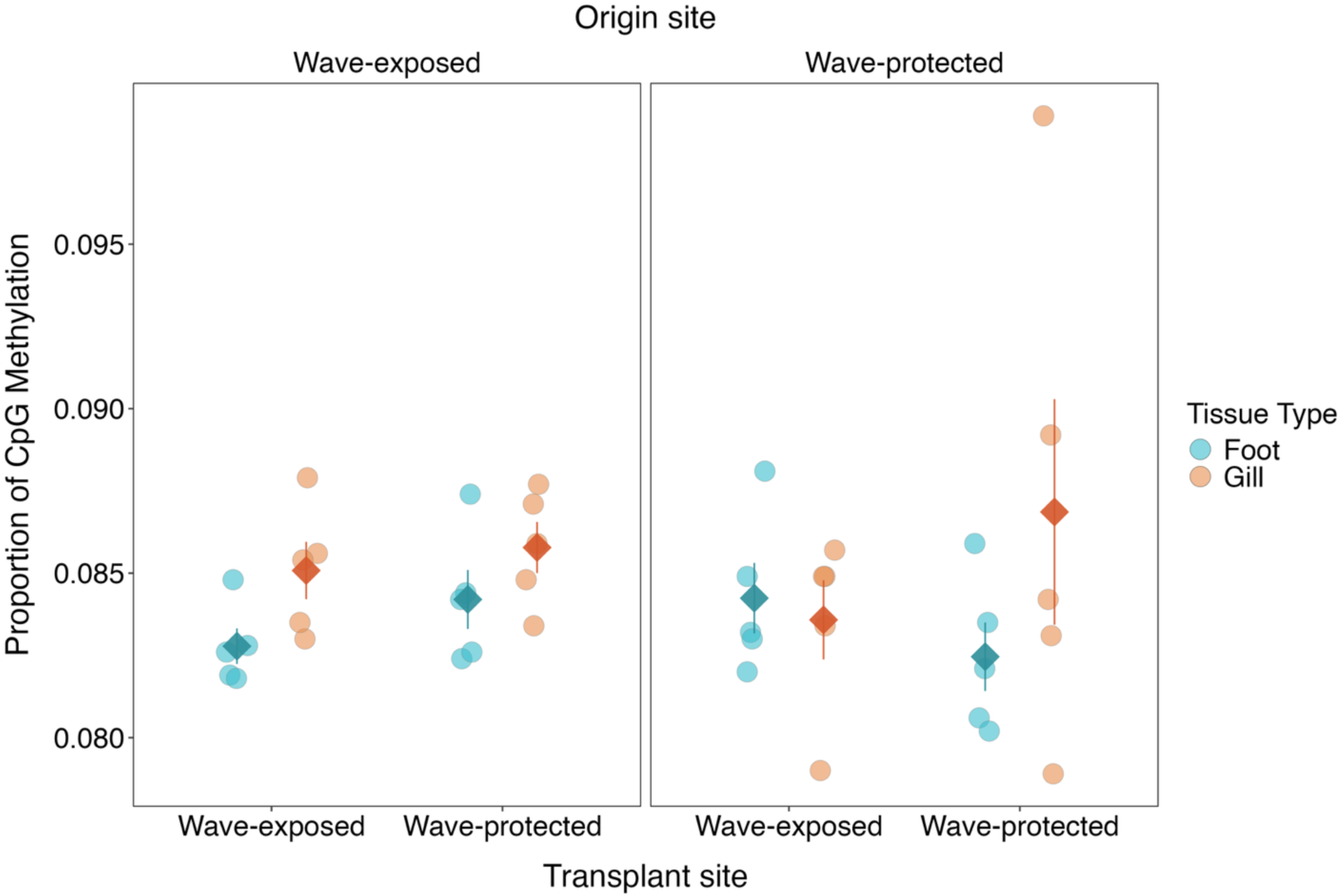
Genome-wide CpG methylation proportion did not significantly differ between gill and foot tissue, and it also was unaffected by origin site (panels) or transplant site (x-axis). Mean genome-wide methylation across CpG loci differed by only 0.19% between foot and gills, by 0.02% between origin sites (wave-protected vs. wave-exposed), and by 0.09% between transplant sites (wave-protected vs. wave-exposed). Outlier samples were removed prior to this analysis, after which only the 5 samples per treatment with the largest library sizes were retained. Each point represents an individual mussel. Diamonds represent the mean methylation proportion in a treatment, and the whiskers represent ± standard error (SE).

### Differential methylation patterns

In the 40 mussel tissue samples selected for all downstream analyses, an average of 65.4 ± 4.17% of reads mapped to the bisulfite-converted *M. californianus* reference genome. An average of 36.5 ± 2.86% of reads aligned uniquely (Table S2). For foot samples, a total of 1,946,085 unique CpG loci from the merged top and bottom strands were identified. After filtering for sites with at least 3x coverage in at least 14 samples, 77,434 (3.98%) CpG sites remained with an average coverage of 13.08 ± 8.84 reads per CpG (Table S2). For gill samples, a total of 2,485,629 unique CpG loci were identified. After applying the same filtering threshold, 76,636 (3.1%) CpG sites remained with an average coverage of 11.15 ± 6.13 reads per CpG (Table S2).

Separate *edgeR* models of DM were fitted to methylation data from foot and gill, using origin and transplant site as fixed effects and shell length as a covariate. Initial versions of these models that also included an origin x transplant interaction term caused overfitting, evidenced by abnormally high absolute log2FC values for many CpGs of both tissues. Additionally, the interaction term explained little variation in methylation in either tissue, with only a small number of significant DM CpGs associated with the origin x transplant interaction effect. (Supplemental Materials; Figures S3-S4). To report conservative estimates of DM, the models of DM described here do not include an origin x transplant interaction.

#### Foot

Using an FDR-adjusted alpha value of 0.05, foot samples for mussels of wave-protected origin exhibited 93 DM CpGs relative to those from the exposed origin site (0.12% of all CpGs in filtered foot dataset; Figure 3A, Table 1). Of these 93 loci, 49 were hypomethylated and 44 were hypermethylated for protected origin mussels (Table S4a). A total of 23 unique genes overlapped with these DM CpGs, including several genes with clusters of intronic CpGs that were DM in the same direction (top five genes Table 2, full list in Table S4a). The genes with the largest numbers of intronic DM CpGs were glycerophosphocholine cholinephosphodiesterase ENPP6-like (4 CpGs) and SID1 transmembrane family member 1-like (3 CpGs). A Wilcoxon signed-rank test showed that DM associated with origin site in foot did not skew toward hyper- or hypo-methylation (p > 0.05). No under- or over-representation of foot DM CpGs was detected across any genomic feature relative to genomic background (Fisher’s exact test & GLM p > 0.05; Table S3c, S3d, Figure 4A).

**Figure 3.**
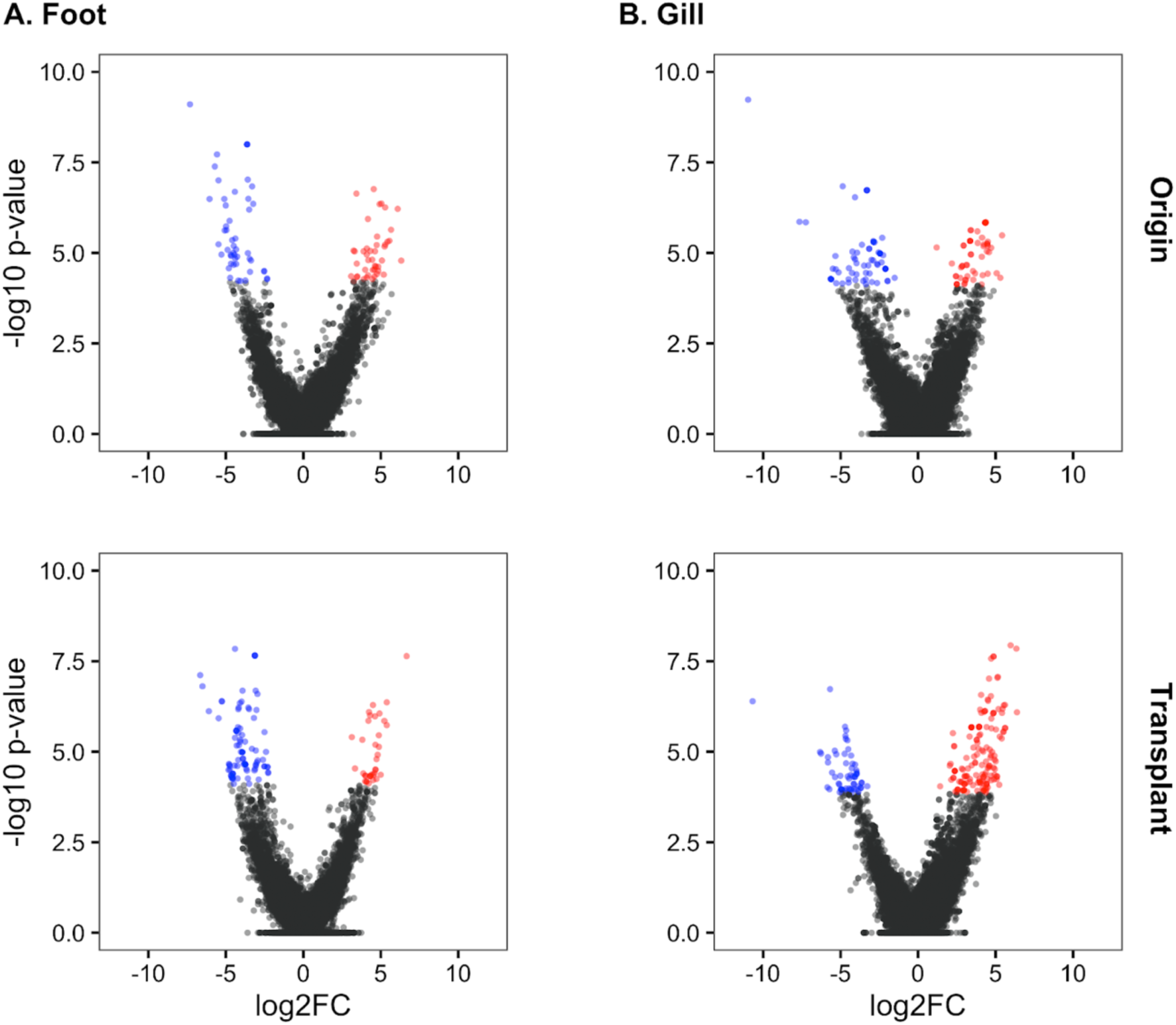
Differentially methylated (DM) CpGs identified for **A**. foot and **B**. gill samples based on origin site (top) and transplant site effect (bottom). Each dot represents a CpG locus. Significantly hypermethylated (red) and hypomethylated (blue) CpGs are indicated, based on an FDR-adjusted alpha-value of <0.05. Exposed sites were used as the reference level for each comparison.

**Figure 4.**
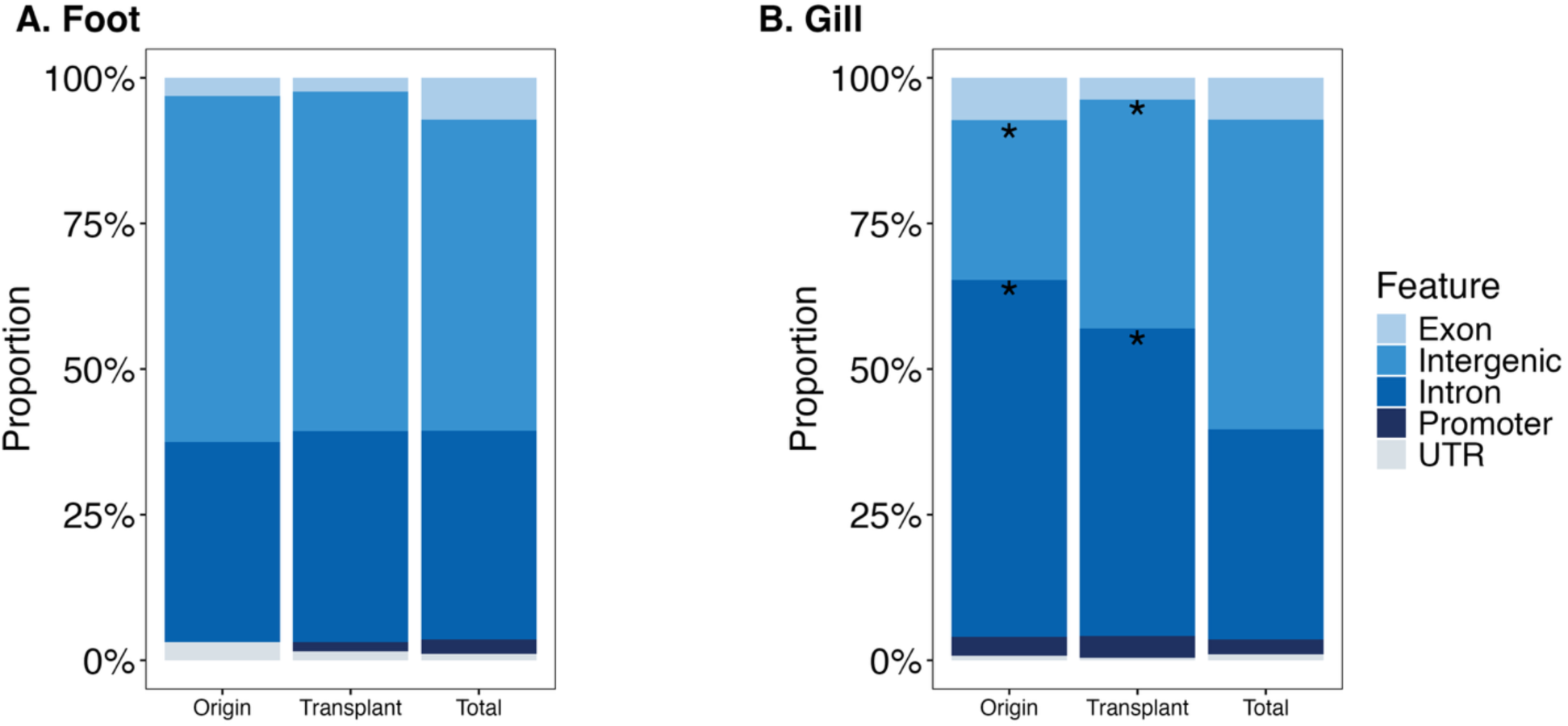
Proportions of differentially methylated (DM) CpGs associated with origin and transplant site effects compared to the background distribution of CpGs within each genomic feature in **A.** foot and **B.** gill. Asterisks in panel B indicate significant enrichment or under-representation of DM CpGs relative to the genomic backgrounds. In gill, DM CpGs were significantly enriched in introns and significantly under-represented in intergenic regions for both origin and transplant effects.

**Table 1.** Distribution of differentially methylated (DM) CpGs across all feature types per tissue associated with origin site, transplant site, and length effects. Seven foot and ten gill DM CpGs were annotated with multiple genomic features. Rows in bold represent the total unique number of CpGs for each tissue and for each effect tested.

| Tissue | Feature | Origin DM | Transplant DM | Length DM | Total CpGs |
| --- | --- | --- | --- | --- | --- |
| Foot | Exon | 3 | 3 | 7 | 5694 |
|  | Intron | 33 | 46 | 63 | 28509 |
|  | Promoter | 0 | 2 | 9 | 1976 |
|  | Intergenic | 57 | 74 | 70 | 42470 |
|  | UTR | 3 | 2 | 0 | 862 |
|  | **Unique** | **93** | **124** | **148** | **77434** |
| Gills | Exon | 9 | 8 | 10 | 5645 |
|  | Intron | 76 | 114 | 84 | 28378 |
|  | Promoter | 4 | 8 | 2 | 1990 |
|  | Intergenic | 34 | 85 | 53 | 41859 |
|  | UTR | 1 | 1 | 1 | 801 |
|  | **Unique** | **121** | **211** | **148** | **76636** |

**Table 2.**
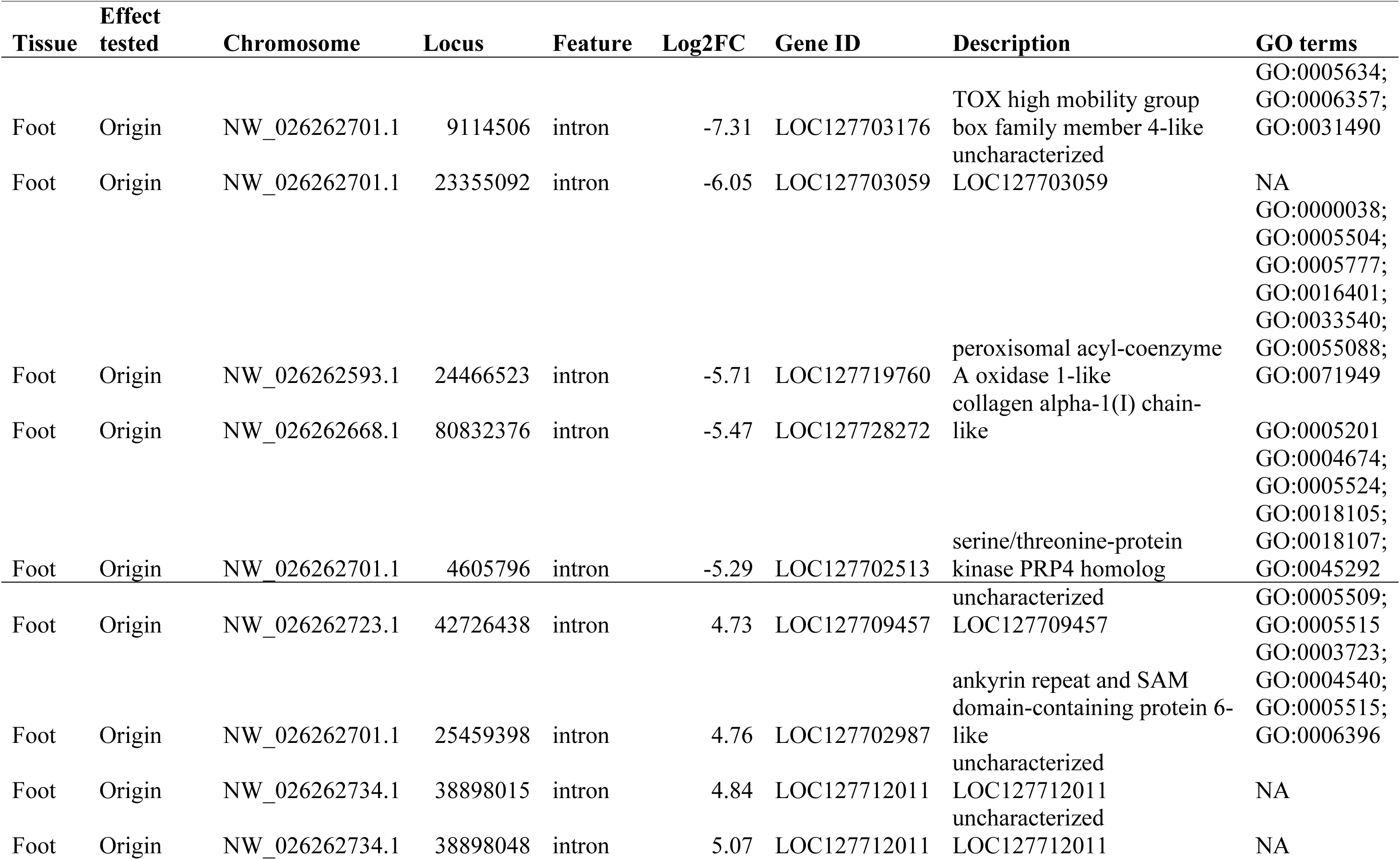
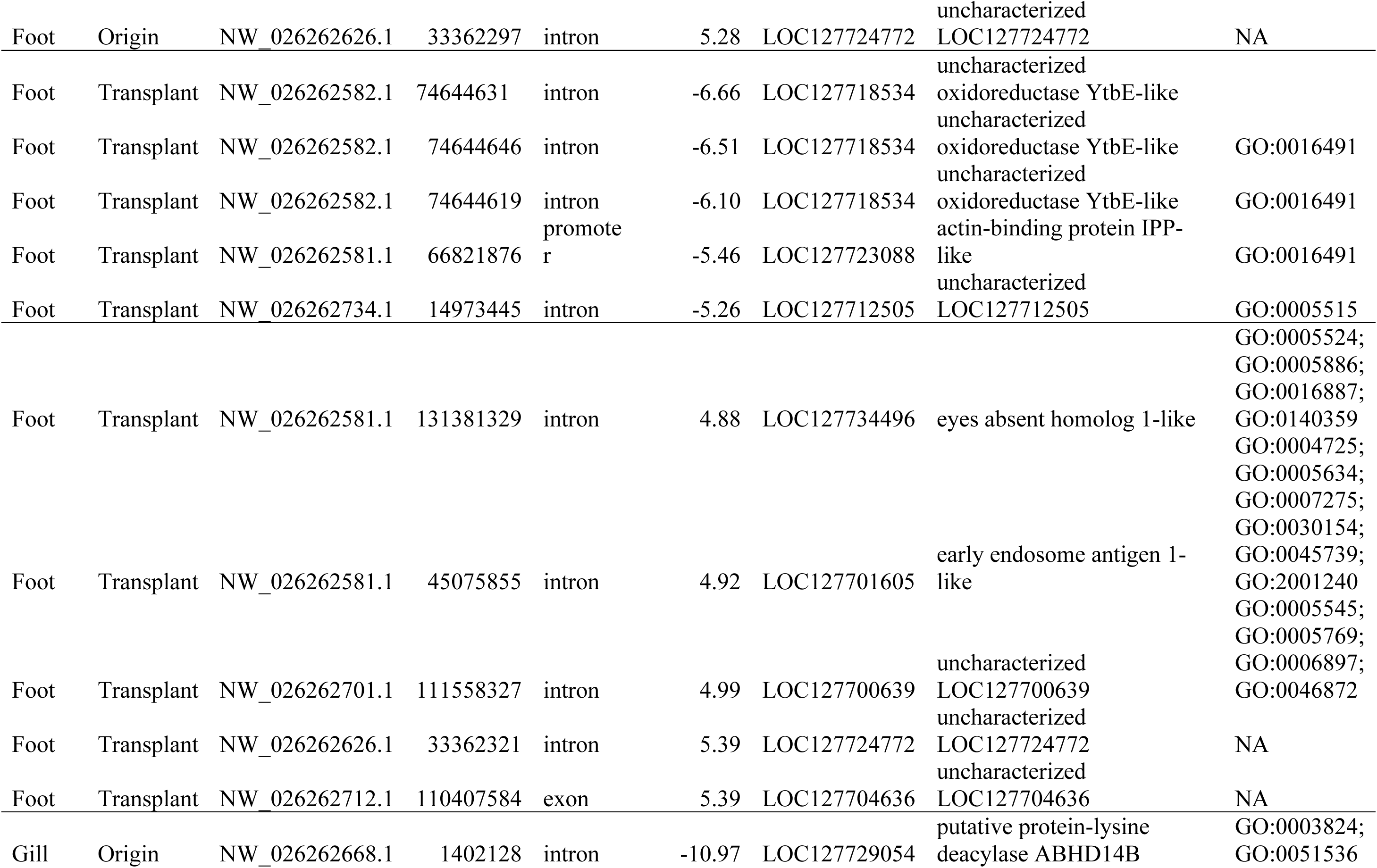
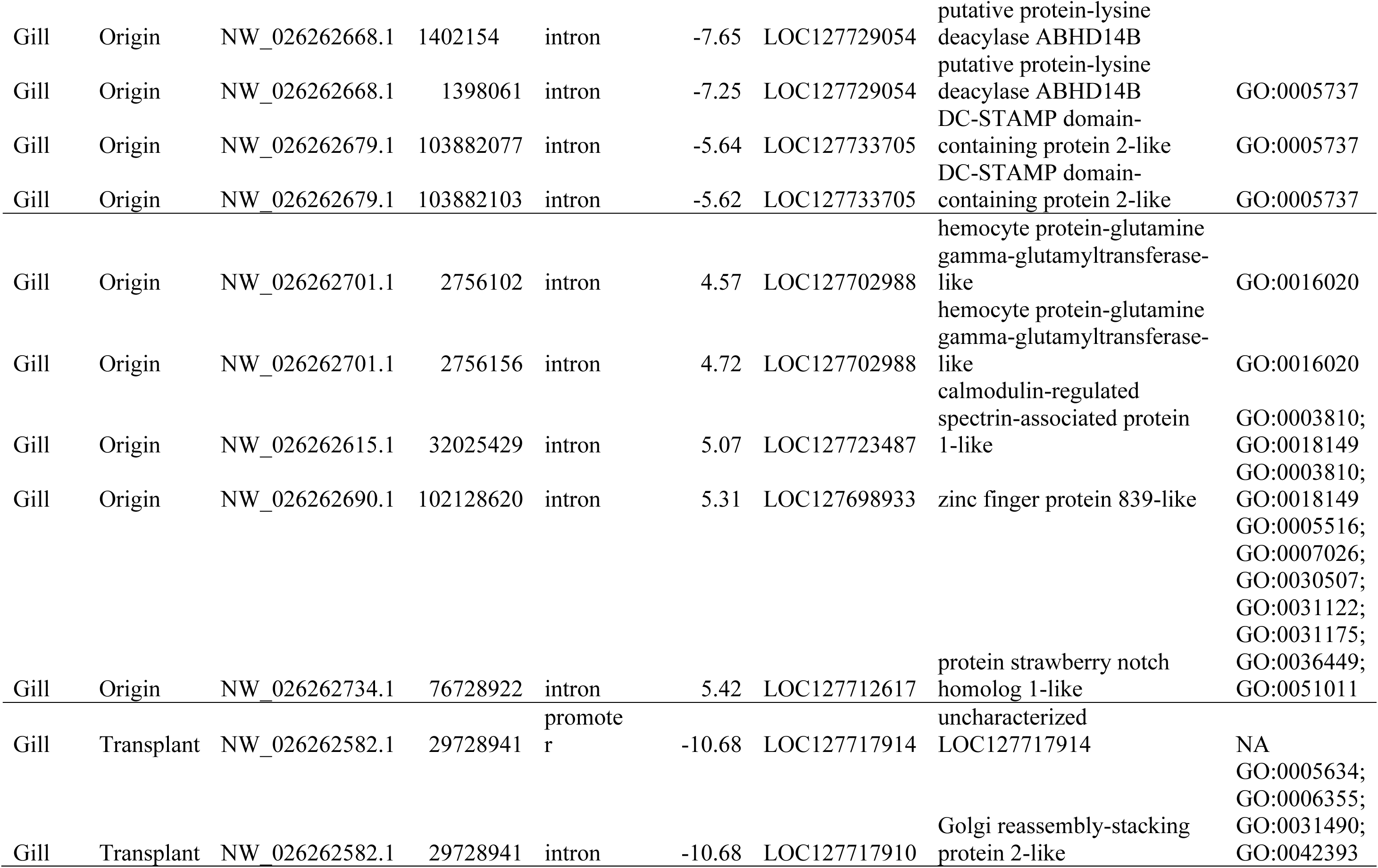
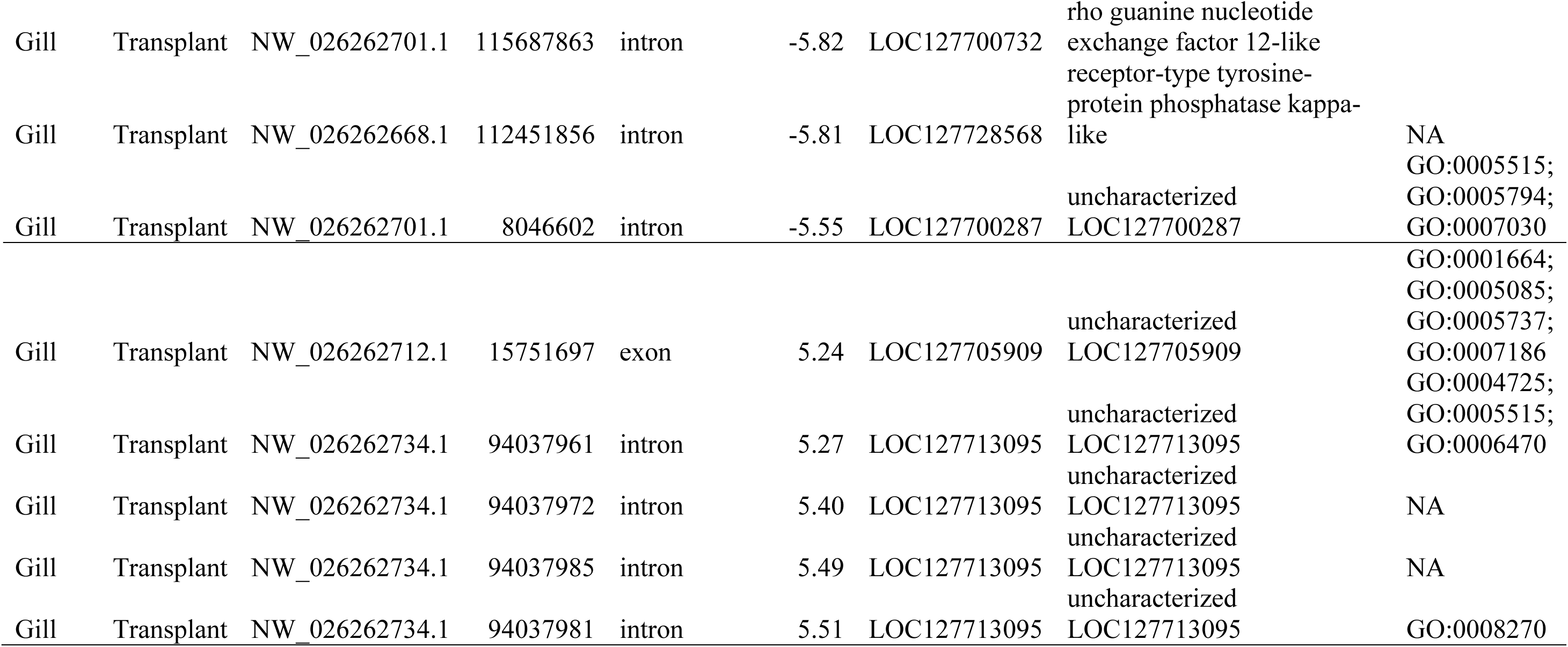
The five most hyper- or hypomethylated gene body CpG loci in each mussel tissue for the effects of origin and transplant site.

In response to transplantation to wave-protected sites, 124 DM CpG sites (0.16% of all foot CpG sites; Figure 3A, Table 1) were identified in foot (88 hypomethylated and 36 hypermethylated loci). These loci overlapped with 28 unique genes, several of which exhibited clusters of intronic DM CpGs that were regulated in the same direction (top five genes Table 2, full list in Table S4b). Among the DM genes with annotations, the largest such clusters of intronic DM CpGs occurred in glycerophosphocholine cholinephosphodiesterase ENPP6-like (4 CpGs), guanine deaminase-like (3 CpGs), probable E3 ubiquitin-protein ligase HERC4 (3 CpGs), and uncharacterized oxidoreductase YtbE-like (3 CpGs). A Wilcoxon signed-rank test showed that transplantation to the wave-protected site resulted in DM in foot that was significantly skewed toward hypomethylation (p < 0.001, Wilcoxon effect size: r = 0.034). Here, r is a measure of skewness calculated as the z-score distance of a distribution’s mean from zero, divided by the square root of the sample size. No genomic feature contained an under- or over-representation of DM CpGs compared to the background across the genome (Fisher’s exact tests & GLM, p > 0.05; Table S3c & 3d, Figure 4A).

The mean log2FC values for all DM CpGs differed significantly between the origin and transplant site effects, although both indicated a predominance of hypomethylation in wave-protected mussels (mean origin effect: -0.15, mean transplant effect: -1.46; LM, p = 0.019). The strength of DM was stronger in response to the origin site than to the transplant site (absolute log2FC: mean origin effect: 4.40, mean transplant effect: 4.07; LM, p = 0.02). Among all DM CpGs, twenty-four unique DM CpGs in foot were shared between the two effects (Figure 5A, highlighted rows in Table S4a, S4b), 19 of which were DM in the same direction. These 24 DM CpGs were divided between intergenic (14) and intronic (10) features, the latter of which overlapped with six unique genes. Notably, the cluster of 4 intronic DM CpGs in glycerophosphocholine cholinephosphodiesterase ENPP6-like were DM in opposite directions for the origin and transplant effects.

**Figure 5.**
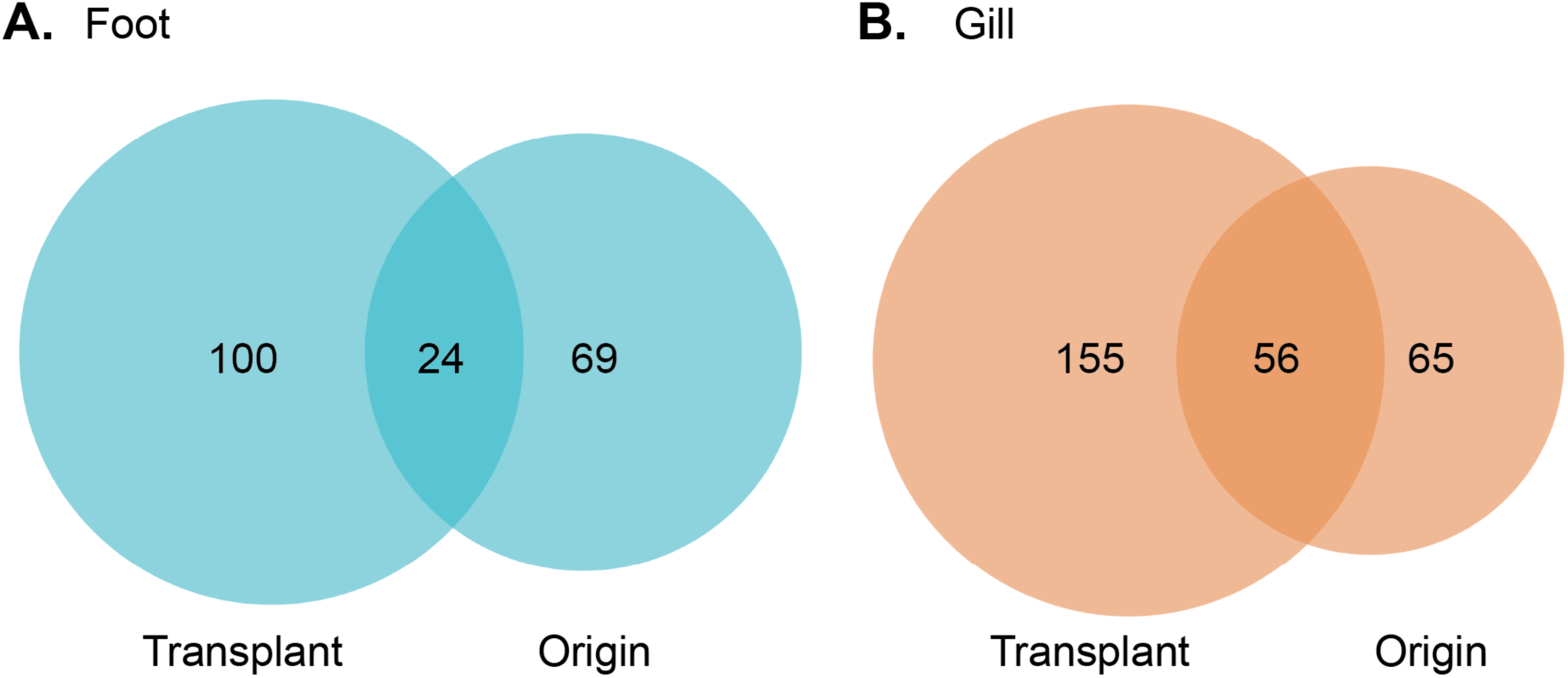
The sets of differentially methylated (DM) CpGs in juvenile mussels were largely distinct between tissues as well as between origin and transplant effects. The overlaps between circles indicate DM CpGs that were shared between effects in either **A.** foot (blue) or **B.** gill (orange). Of 63,586 total CpGs shared between the tissues, only one was DM for the origin effect in both tissues, and there was no overlap in DM CpGs between tissues for the transplant effect.

There was a total of 148 significant DM CpGs in foot associated with shell length (70 negatively correlated and 78 positively correlated; Table 1, Table S4c). These DM CpGs overlapped with 41 genes.

#### Gill

Gill in mussels of wave-protected origin exhibited 121 DM CpG sites compared to mussels from the exposed origin site (0.16% of all gill CpG sites; Figure 3B, Table 1). The 64 hypomethylated and 57 hypermethylated DM CpGs overlapped with 45 unique genes (top five genes Table 2; full list in Table S5a). As in foot, within some of these genes, there were clusters of intronic DM CpGs that were regulated by origin site in the same direction; these included protein SSUH2 homolog (7 CpGs), mucin-22-like (7 CpGs), putative protein-lysine deacylase ABHD14B (5 CpGs), and transient receptor potential cation channel subfamily M member-like 2 (5 CpGs). A Wilcoxon signed-rank test showed that DM associated with wave-protected origin was significantly skewed toward hypomethylation (p < 0.001; Wilcoxon effect size: r = 0.05). DM CpGs in gill associated with origin effect were significantly enriched in introns and significantly underrepresented in intergenic regions compared to genomic background (Fisher’s exact test, p < 0.001 for each, Table S3c, Figure 4B). Intergenic regions were under-enriched in DM CpGs compared to introns (binomial GLM, p < 0.05; Table S3d, Figure 4B).

The largest number of DM CpGs of any comparison occurred in gill in response to transplant to the wave-protected site (211 CpGs, 0.28% of analyzed CpG sites; Figure 3B, Table 1). This list included 1.7 times the number of DM sites associated with the transplant effect in foot. This set of DM CpGs included 59 that were hypomethylated and 152 that were hypermethylated in the wave-protected transplant site (Table S5b). These DM CpGs overlapped with 62 different genes (top five genes Table 2, full list in Table S5b), once again including numerous clusters of intronic DM CpGs. The largest such intronic clusters within annotated genes were associated with protein SSUH2 homolog (7 CpGs), mucin-22-like (7 CpGs), nucleolar protein dao-5-like (7 CpGs), transient receptor potential cation channel subfamily M member-like 2 (5 CpGs), and receptor-type tyrosine-protein phosphatase kappa-like (5 CpGs). Contrary to all other effect-by-tissue combinations, gill DM caused by transplantation to the wave-protected site was significantly skewed towards hypermethylation (Wilcoxon signed rank test, p < 0.001; effect size: r = 0.07). Transplant DM CpGs in gill were significantly enriched in introns and underrepresented in intergenic regions compared to the genomic background (Fisher’s exact test, p < 0.001 for each, Table S3c, Figure 4B). Introns were more likely to contain DM CpGs compared to both exons and intergenic regions (binomial GLM, p < 0.05 for each; Table S3d, Figure 4B).

The strength and directionality of DM in all DM CpGs differed significantly between the origin and transplant site effects. On average, DM changes associated with transplant site were stronger and were in the opposite direction compared to those associated with origin site (absolute log2FC: mean origin effect = 3.70, mean transplant effect = 4.20; LM, p < 0.001 for both). Among all DM CpGs, a total of 56 unique DM gill CpGs were shared between the two effects (Figure 5B, highlighted rows in Table S5a, S5b). The distribution of shared DM CpGs across genomic features in gill differed from foot, with 16 intergenic, 35 in introns, 2 in exons, 3 in promoters, and 1 in UTR that overlaps with an exon. The shared intragenic DM CpGs in gill overlapped with 14 unique genes. Also, unlike foot, 45 of the 56 shared sites being regulated in opposite directions for origin and transplant.

Finally, the effect of shell length was associated with 148 gill DM CpGs, of which 83 were negatively correlated and 65 were positively correlated (Table 1, Table S5c). These DMs overlapped with 46 unique genes.

#### Tissue-specificity of differential methylation induced by origin and transplant site

Mussels’ foot and gill CpG methylation patterns responded very differently to the origin and transplant effects. For individual DM CpGs, there was minimal overlap between the two mussel tissues. In response to origin site effect, only one intergenic DM CpG was shared by both tissues (NW_026262690.1:69391943), and no DM CpGs were shared between tissues in response to transplant effect. The distribution of DM CpGs across genomic features in gill also significantly differed from that in foot. For both origin and transplant effects, intergenic regions were enriched and introns were depleted for DM in the foot as compared to gill (Fisher’s exact tests, p < 0.01 for each; Table S3c). Additionally, we assessed whether the strength and directionality of DM differed between the two tissue types. For mussels from the wave-protected origin site, the mean log2FC values for each tissue indicated hypomethylation and were not significantly different (foot: -0.15, gill: -0.27; LM, p = 0.83); however, the overall strength of DM for the origin effect was significantly greater in foot (absolute value of log2FC: foot: 4.40; gill: 3.69, p < 0.0001). The tissues differed in their response to transplantation to wave-protected sites (LM, p < 0.0001); foot was more likely to exhibit hypomethylation (mean log2FC: -1.46), whereas gill on average exhibited hypermethylation (mean log2FC: 1.56). Unlike for the origin effect, for transplant there was no significant difference between tissues in the overall strength of DM (absolute value of log2FC: foot: 4.07, gill: 4.20; LM, p = 0.30). Lastly, we assessed the relationships between the log2FC values for the two tissues (using all shared CpG sites in the dataset). For both origin and transplant effects, there existed a significant but very weak positive correlation between the log2FC values in the two tissues (Pearson’s correlation, origin r = 0.11, transplant r = 0.069, p < 0.001 for each; Figure S5).

### Gene Ontology (GO) Enrichment Analysis

For all treatment groups in both tissues, no GO terms were enriched for the sets of DM genes after FDR correction using an alpha value < 0.05. Summaries of the GO enrichment analysis are available in Supplemental Materials (Table S6).

## Discussion

Methylation of DNA at CpG loci, one of the most widely studied epigenetic mechanisms, is known to contribute to gene regulation underlying plastic responses to environmental stressors (Bogan & Yi, 2024; Duncan et al., 2014; Eirin-Lopez & Putnam, 2019; Sun et al., 2022). To understand how different *in situ* habitat conditions (cool, wave-exposed vs. warm, wave-protected intertidal sites) influence DNA methylation in juveniles of *M. californianus*, a sessile intertidal invertebrate, we performed RRBS on gill and foot samples from mussels reciprocally conditioned to different intertidal wave-exposure regimes (Gleason et al., 2018). The use of a reciprocal transplant design allowed for the identification of plastic DNA methylation responses to the environment in two distinct tissue types, which has rarely been possible in ecological epigenetics studies.

The plastic responses identified in this study were subtle and did not involve wholesale shifts in CpG methylation levels. Indeed, multivariate patterns of methylation across all CpG loci exhibited no evidence of clustering of samples by tissue type, origin site, or transplant treatment. This lack of multivariate clustering by treatment group aligns with findings from a study on the eastern oyster (*Crassostrea virginica*), where gonad samples exposed to control or elevated pCO_2_ showed no separation when considering all CpG loci (Venkataraman et al., 2020). Furthermore, we did not observe any significant treatment effects on genome-wide methylation levels in mussel foot or gill. Genome-wide DNA methylation levels in the California mussel were consistently low in both tissues (<10%, Figure 2) and comparable to values of ∼15-35% observed in other molluscs (Roberto et al., 2021; Zhang et al., 2020). The lack of significant changes in mussels’ genome-wide methylation levels contrasts previous studies that documented a significant decrease in global DNA methylation in the eastern oysters (*C. virginica*) under ocean acidification (Downey-Wall et al., 2020) and a significant increase in global DNA methylation in the sea cucumber (*Apostichopus japonicus*) in response to heat stress (Chang et al., 2023).

Nonetheless, the one-month reciprocal transplant experiment led to a number of significant changes in methylation of specific CpGs in foot and gill that generally supported our predictions. Both the transplant environment (wave-exposed or wave-protected) and the origin environment influenced DNA methylation, but a larger number of DM CpGs were associated with the recently experienced transplant conditions. As predicted, distinct sets of DM CpGs (and the genes they resided in) were found in foot and gill in response to the origin and transplant environments. Several of the DM intragenic CpGs were in genes with links to cellular stress responses, generating mechanistic hypotheses for the enhanced heat tolerance observed among mussels transplanted to the wave-protected site (Gleason et al., 2018). Overall, these results demonstrate that environmental effects on DNA methylation are evident even at small spatial scales across the intertidal zone, and that different tissues exhibit distinct, plastic epigenetic responses to microhabitat conditions. As a consequence, the potential organismal implications of environment-methylation associations derived only from single tissues should be assessed cautiously.

### Tissue-specific changes in DNA methylation

Although transplantation between wave-exposed and wave-protected sites did not lead to genome-wide changes in CpG methylation in *M. californianus*, we identified several hundred tissue-specific DM CpGs associated with the effects of microhabitats. As predicted, both the number and the distribution of DM CpGs differed between tissues. The number of DM CpGs associated with transplant site effect in gill was 1.7x higher than those in foot. The directionality of DM in response to the transplant environment also differed between the two tissue types, whereas the strength of DM associated with the origin site effect was greater in foot compared to gill. Furthermore, for both origin and transplant effects, there existed only a very weak positive correlation in the log2FC values of all shared CpG loci between the two tissue types. In fact, only 1 DM CpG was shared between foot and gill for any comparison, and the sets of genes containing DM CpGs in each tissue also had little overlap.

These results are consistent with distinct patterns of epigenomic plasticity between the two tissue types, potentially reflecting their distinct biological functions. For example, greater epigenomic plasticity may be found in gills due to their numerous physiological roles (Dahlhoff et al., 2002; Thompson & Bayne, 1972). In the laboratory, acute heat stress induced significant changes in over 1,500 transcripts in gills of adults *M. californianus*, including genes whose products are involved in protein folding and proteolysis (Lockwood & Somero, 2011); similar functions were represented among the differentially methylated genes in our findings. In a field study, adult *M. californianus* outplanted to a hot, high-intertidal site exhibited differential expression in gills for hundreds of genes, including some involved in chromatin structural organization (e.g., *histone H1-delta like* and *chromobox protein homolog 1-like*) (Gleason et al., 2023). These gene products might also play a role in DNA methylation (Geeven et al., 2015; Wang et al., 2023), suggesting that gill epigenomic plasticity could contribute to mussels’ acclimatization to the environment at multiple life history stages. In contrast, responses to heat stress in foot have been less studied in mussels and were previously known to increase the expression of proteins involved in byssal thread formation (Li et al., 2020).

Our results are consistent with past studies in molluscs, which provided evidence of tissue-specific methylation responses to biological and environmental stressors. For example, tissue-specific methylation changes were found in the immune genes of the gills and mantles of the Pacific oyster (*Magallana gigas*) after a Pacific Oyster Mortality Syndrome infection, potentially reflecting the different roles the two tissue types play in the immune response (Valdivieso et al., 2025). Differences in DNA methylation were also observed in the sperm and egg tissues of the eastern oyster (*Crassostrea virginica*) after exposure to elevated pCO_2_; methylation responses in sperm were more pronounced and showed an opposing relationship with gene expression when compared to eggs (Venkataraman, Huffmyer, et al., 2024).

Interestingly, tissues from the same individual clustered more closely together in multi-dimensional PC space than those collected from different individuals. Differences in body size (here, shell length) explained a small portion of this pattern. Other factors not included in our analysis, such as genetic variation, likely also contributed. For example, 27% of interindividual variation in methylation in the Olympia oyster (*Ostera lurida*) was explained by genetic distance (Silliman et al., 2023). Although *M. californianus* populations exhibit little genetic evidence of local adaptation throughout their range in the northeast Pacific (Addison et al., 2008; Levinton & Suchanek, 1978), each population can exhibit considerable genetic diversity (Ort & Pogson, 2007). Future epigenomic research in *M. californianus* and other genetically diverse, widely-dispersing organisms should account for genetic effects on DNA methylation (Bogan & Yi, 2024). An alternative, but less likely, explanation for the lack of sample clustering by tissue is that epigenomic divergence between foot and gills is limited.

### Effects of intertidal wave-exposure conditions on DNA methylation in stress-response genes

The previously documented increase in whole-organism heat tolerance of juvenile mussels transplanted to the wave-protected site strongly indicates that they were able to compensate for more intense stressors (Gleason et al., 2018). The average daily range of mussel body temperatures was three times greater at the protected site compared to the exposed site during the transplant experiment (8.1 °C vs. 2.7 °C; Gleason et al., 2018). More frequent episodes of high body temperature during low tide likely caused macromolecular damage and oxidative stress in the juvenile mussels at that site (Halliwell & Gutteridge, 2015; Jimenez et al., 2016; Stillman et al., 2025). Accordingly, transplant site was associated with a larger number of DM CpGs than origin site in both tissues, and several of the methylation responses associated with transplant site mapped to genes involved in functions related to combating abiotic stress. For example, several transplant DM genes have putative roles in proteostasis (chaperone and protein degradation; including protein SSUH2 homolog, endoplasmic reticulum resident protein 44-like, E3 ubiquitin-protein ligase DZIP3-like), DNA repair (e.g., TOX high mobility group box family member 4-like), regulation of transcription (including RNA processing), maintenance of Golgi apparatus structure, cytoskeletal organization (including cilia structure and cell-cell adhesion), lipid metabolism, and signaling pathways. All of these processes are relevant to the conserved cellular stress response (Kültz, 2003, 2005).

These responses to transplantation in *M. californianus* align with a previous study by Clark et al. (2018), which observed that reciprocal transplantation of Antarctic limpets (*Nacella concinna*) between intertidal and subtidal zones resulted in significantly different methylation patterns. However, the amplified fragment length polymorphism technique used to assess DNA methylation in that study prevented resolution of specific genomic features. The current results clarify that multiple tissue types, gene functional categories, and genomic features (exons, introns, intergenic regions, promoters, and UTRs) exhibit plastic DNA methylation responses to environmental variation within the intertidal zone.

We also predicted that more mussel stress-response genes would be differentially methylated in response to manipulation of their intertidal wave-exposure regime in gills than in foot, due to gills’ multiple physiological roles and their known susceptibility to environmental stressors. Tissue-specific vulnerability was observed in the mussel *M. edulis* and the clam *Dosinia lupinus*, which accumulated different levels of protein carbonyls across four tissues (digestive gland, mantle, adductor muscle, and foot) following an exposure to oxidative stress (Walker et al., 2000). As noted above, both juvenile mussel tissues exhibited differential methylation within annotated genes associated with various aspects of the cellular stress response. However, there were no significantly enriched GO terms associated with the sets of DM genes for either tissue after correction for multiple testing. RRBS often lacks the power to detect functional enrichment (Hu et al., 2021; Seiler Vellame et al., 2021; Zhou et al., 2016), given that this approach only identifies a limited subset of CpGs and the genes that contain them. Furthermore, many of the DM genes remain unannotated or have not been characterized functionally in mussels. Ultimately, our prediction is difficult to evaluate quantitatively.

Despite these limitations, a qualitative assessment of the putative functions of the DM genes highlights several differences between tissues and/or treatment effects that warrant further exploration in the context of mussels’ physiological plasticity. First, gill transplant effect was associated with a relatively large number of DM genes with putative roles in the regulation of transcription and RNA processing (including EEF1A lysine methyltransferase 4-like; pre-mRNA-splicing factor RBM22-like; U2 snRNP-associated SURP motif-containing protein-like; integrator complex subunit 3-like) as well as genes associated with ribosomes (nucleolar protein dao-5-like; 60S acidic ribosomal protein P2-like; 60S ribosomal protein L5-like). In contrast, these functions did not feature among foot transplant DM genes. Regulation of gene expression plays an important role in the acclimation of organisms to abiotic stress (including thermal stress) by allowing them to modify their phenotypes to enhance their cellular stress resistance (Kültz, 2005; Somero, 2010). Similarly, gill transplant effect was associated with a considerable number of DM genes involved in cell structure and cell-cell adhesion. Among these were several genes that contribute to cilia formation (including CBY1-interacting BAR domain-containing protein 1-like; centrosomal protein of 128 kDa-like) or to mucins and glycans in the extracellular matrix (mucin-22-like; beta-1,2C3-galactosyl-O-glycosyl-glycoprotein beta-1,2C6-N-acetylglucosaminyltransferase-like; L-fucose kinase-like). The numerous cilia of mussel gills are essential for feeding, and past work has hypothesized that the cytoskeletal structure of these fine projections might be sensitive to heat stress (Lockwood & Somero, 2011). Extracellular mucins and glycans contribute to the extracellular matrix (Cerullo et al., 2020; Corfield, 2015), and gills could rely on these materials to maintain the integrity of their organism-environment interface in the face of temperature stress (Chen et al., 2019; Pales Espinosa et al., 2016). Effects of thermal stress on the integrity of the gill barrier might also relate to the observation of gill DM genes that are putatively involved in immune responses (basic immunoglobulin-like variable motif-containing protein; CD109 antigen-like); other studies have observed regulation of immunity genes in response to heat stress (Ericson et al., 2024; Lang et al., 2009; Li et al., 2017). In addition, cysteine sulfinic acid decarboxylase-like, which catalyzes a key step in taurine biosynthesis, was also differentially methylated in response to transplant (and origin) only in gills. Taurine levels in gills have previously been shown to positively correlate with thermal history in adult mussels (Gleason et al., 2017), and taurine may function as a thermoprotectant (Arakawa & Timasheff, 1985; Huxtable, 1992). In contrast, a larger number of transplant DM genes were putatively associated with DNA repair processes in foot than in gills, which might suggest different levels of DNA damage caused by thermal stress in the two tissue types (González-Riopedre et al., 2007). Finally, in foot the gene for glycerophosphocholine cholinephosphodiesterase ENPP6-like, which liberates choline and/or processes degradation products of the membrane constituent phyosphatydylcholine, was differentially methylated for both transplant and origin effects. PC is an important component of the cell membranes of animals (Hochachka & Somero, 2002; Nguyen et al., 2020; Somero et al., 2016; van der Veen et al., 2017). Mussels are known to rapidly remodel membrane order under changing conditions (via homeoviscous acclimation; Williams & Somero, 1996), and it is possible that shifts in PC content in foot membranes also contribute to this process and to sustained organismal function in the face of thermal extremes. In each case, further physiological studies are needed to evaluate these hypothesized contributions to thermal acclimatization in juvenile mussels.

### Effect of body size on DNA methylation

In the original reciprocal transplant experiment, Gleason et al. (2018) observed a trade-off between thermal tolerance and growth rate among juvenile *M. californianus*. Juvenile mussels transplanted to the protected sites grew more slowly than those at the exposed sites, possibly due to energetic constraints (Gleason et al., 2018). Fitzgerald-Dehoog et al. (2012) also showed that adult *M. californianus* exposed to thermal stress reduced shell growth. DNA methylation can have an impact on organismal growth patterns. For example, DNA methylation was shown to be involved in muscle development and growth in fishes (Huang et al., 2018; Kho & Ruzzante, 2024; Pan et al., 2021). Epigenetic mechanisms also can underlie growth patterns in the Pacific Oyster (Tan et al., 2022). Our multivariate PCA analysis revealed that 37% of the variation in PC1 could be explained by shell length of *M. californianus*; however, PC1 only explained 3.75% of the total variation in CpG methylation, and PC2 was not correlated with shell length. We also observed numerous DM CpGs associated with the effect of shell length in both tissues. These relationships suggest that CpG DNA methylation may play a role in mussel development (Haidar et al., 2024; Tan et al., 2022; Vogt, 2022), perhaps independent of environmental conditions. Additional studies with larger sample sizes taken over the course of mussels’ development would be needed to more fully evaluate these potential mechanisms. Recently, whole genome bisulfite sequencing of *M. californianus* revealed that CpG methylation associates with age (Hiebert et al., 2025). Because *M. californianus* age and shell length correlate, it is possible that shell length-associated methylation detected in our study was caused by mussel age rather than length. Regardless, controlling for the impact of length or age on CpG methylation was critical for measuring differential methylation between environments.

### Functional implications of DNA methylation changes

As noted above, the true implications of shifts in CpG methylation for cellular and organismal physiology of juvenile mussels await further evaluation. The complex relationship between methylation changes and gene expression (i.e., transcript and protein levels) also warrants caution in interpretation. In general, associations between environmentally induced changes in DNA methylation and gene expression tend to be weak in invertebrates (Bogan & Yi, 2024; Dixon & Matz, 2022; Duncan et al., 2022; Oldroyd & Yagound, 2021). In the few studies in molluscs, environmentally induced differential DNA methylation and differential gene expression lacked correlation when considered on a genome-wide scale (Downey-Wall et al., 2020; Johnson et al., 2022). Of course, such global analyses leave open the possibility that individual genes might exhibit stronger correlations. In addition, alterations in DNA methylation may have effects on gene expression that do not manifest as differences in transcript levels. In some instances, differential methylation may instead stabilize transcription levels, thereby promoting transcriptional homeostasis, or it can modify chromatin structure that alters accessibility of genes to the transcriptional machinery and regulate transcription (Bogan & Yi, 2024; Li et al., 2018; Ng & Adrian, 1999).

In this context, one notable feature of our results is the enrichment of DM CpGs within specific genomic features, some of which may be more likely to impact transcript levels in invertebrates. DNA methylation levels at CpG loci in invertebrates are known to be lower genome-wide, but they tend to be concentrated in gene bodies compared to those in vertebrates (Klughammer et al., 2023). Interestingly, in response to both origin and transplant effects in gills, intergenic DM CpGs were shown to be under-represented while DM CpGs were enriched in introns compared to the rest of the genomic background. In both the purple sea urchin (*Strongylocentrotus purpuratus*) and the honeybee (*Apis mellifera carnica*), genes containing introns with high levels of methylation are more likely to exhibit transcript variants consistent with alternative splicing, alternative transcriptional start sites, and/or exon skipping (Bogan et al., 2023; Flores et al., 2012). In response to transplantation to the wave-protected site in both tissue types, we also observed clusters of CpGs within one or a few introns of a single gene that were differentially methylated in the same direction. These genes with clusters of environmentally responsive intragenic methylation sites represent intriguing targets for future study in intertidal organisms.

## Conclusion

This study documented plastic DNA methylation responses to microhabitat changes in gills and foot of juvenile California mussels (*Mytilus californianus*) using a one-month reciprocal transplant between cool, wave-exposed and warm, wave-protected sites. The number of CpG loci differentially methylated due to recent environmental exposure (transplant site) was greater than that for the effect of pre-transplantation environment (origin site) in both tissues. Differential methylation was observed within numerous genes with known or hypothesized roles in abiotic stress tolerance, potentially contributing to the previously documented pattern of increased heat tolerance for juvenile mussels transplanted to the wave-protected site (Gleason et al. 2018). Furthermore, the plastic responses in CpG methylation were almost entirely tissue-specific. Only one DM CpG locus was shared between the two tissues for the effect of origin site wave exposure. Foot and gill tissues also differed in the directionality of methylation changes due to transplantation and in the distribution of those changes across genomic features. This work expands on previous research that examined DNA methylation responses to environmental changes by investigating plastic and tissue-specific changes in DNA methylation *in situ*. While the physiological significance of much of this CpG methylation variation remains unexplained, future research should aim to link these DNA methylation responses with genetic diversity, transcriptomic data, and physiological metrics to better understand its functional implications.

## Author Contributions

R.L.T. and W.W.D. developed the initial research question and performed the RRBS sequencing. Q.C. analyzed the data with support from W.W.D., S.N.B., and J.L.K. Q.C. wrote the initial draft of the manuscript, and all authors contributed to the revisions. All authors approved the final draft of the manuscript for publication.

## Acknowledgements

We thank Lani Gleason for leading the reciprocal transplant experiments for this study. We also thank members of Kelley and Cornejo Labs for feedback. This research used resources of the shared high-performance computing facility Hummingbird at the University of California, Santa Cruz. Support for Hummingbird was provided by the Information Technology Services Division and the Office of Research at the University of California, Santa Cruz. This research was funded by the National Science Foundation (NSF) IOS-1256186 and IOS-2349877 to W.W.D. Q.C. is supported by an NSF Graduate Research Fellowship under Grant No. 2240310. S.N.B. was supported by NSF OPP-2312253 while contributing to this study.

## Data Availability Statement

The datasets generated for this study can be found in online repositories. The raw sequence files can be found in NCBI BioProject PRJNA1369104. Scripts and intermediate outputs can be found at the online GitHub repository: https://github.com/qitingcai/mytilus_methylation_2025

## References

Addison, J. A., Ort, B. S., Mesa, K. A., & Pogson, G. H. (2008). Range-wide genetic homogeneity in the California sea mussel (Mytilus californianus): A comparison of allozymes, nuclear DNA markers, and mitochondrial DNA sequences. Molecular Ecology, 17(19), 4222–4232. 10.1111/j.1365-294X.2008.03905.x

Akalin, A., Franke, V., Vlahoviček, K., Mason, C. E., & Schübeler, D. (2015). genomation: A toolkit to summarize, annotate and visualize genomic intervals. Bioinformatics, 31(7), 1127–1129. 10.1093/bioinformatics/btu775

Almeida, E. A., Bainy, A. C. D., Dafre, A. L., Gomes, O. F., Medeiros, M. H. G., & Di Mascio, P. (2005). Oxidative stress in digestive gland and gill of the brown mussel (Perna perna) exposed to air and re-submersed. Journal of Experimental Marine Biology and Ecology, 318(1), 21–30. 10.1016/j.jembe.2004.12.007

Andrews, S. (2010). *FastQC: A Quality Control Tool for High Throughput Sequence Data* [Computer software]. http://www.bioinformatics.babraham.ac.uk/projects/fastqc/

Arakawa, T., & Timasheff, S. N. (1985). The stabilization of proteins by osmolytes. Biophysical Journal, 47(3), 411–414. 10.1016/S0006-3495(85)83932-1

Bates, D., Mächler, M., Bolker, B., & Walker, S. (2015). Fitting Linear Mixed-Effects Models Using **lme4**. Journal of Statistical Software, 67(1). 10.18637/jss.v067.i01

Benjamini, Y., & Hochberg, Y. (1995). Controlling the False Discovery Rate: A Practical and Powerful Approach to Multiple Testing. Journal of the Royal Statistical Society: Series B (Methodological), 57(1), 289–300. 10.1111/j.2517-6161.1995.tb02031.x

Bogan, S. N., Johns, J., Griffiths, J. S., Davenport, D., Smith, S. J., Schaal, S. M., Downey-Wall, A., Lou, R. N., Lotterhos, K., Guidry, M. E., Rivera, H. E., McGirr, J. A., Puritz, J. B., Roberts, S. B., & Silliman, K. (2023). A dynamic web resource for robust and reproducible genomics in nonmodel species: Marineomics.io. Methods in Ecology and Evolution, 14(11), 2709–2716. 10.1111/2041-210X.14219

Bogan, S. N., Johnson, K. M., & Hofmann, G. E. (2020). Changes in Genome-Wide Methylation and Gene Expression in Response to Future pCO2 Extremes in the Antarctic Pteropod Limacina helicina antarctica. Frontiers in Marine Science, 6. 10.3389/fmars.2019.00788

Bogan, S. N., Strader, M. E., & Hofmann, G. E. (2023). Associations between DNA methylation and gene regulation depend on chromatin accessibility during transgenerational plasticity. BMC Biology, 21(1), 149. 10.1186/s12915-023-01645-8

Bogan, S. N., & Yi, S. V. (2024). Potential Role of DNA Methylation as a Driver of Plastic Responses to the Environment Across Cells, Organisms, and Populations. Genome Biology and Evolution, 16(2), evae022. 10.1093/gbe/evae022

Bossdorf, O., Richards, C. L., & Pigliucci, M. (2008). Epigenetics for ecologists. Ecology Letters, 11(2), 106–115. 10.1111/j.1461-0248.2007.01130.x

Brooks, M., E., Kristensen, K., Benthem, K., J., van, Magnusson, A., Berg, C., W., Nielsen, A., Skaug, H., J., Mächler, M., & Bolker, B., M. (2017). glmmTMB Balances Speed and Flexibility Among Packages for Zero-inflated Generalized Linear Mixed Modeling. The R Journal, 9(2), 378. 10.32614/RJ-2017-066

Buckley, B. A., Owen, M.-E., & Hofmann, G. E. (2001). Adjusting the thermostat: The threshold induction temperature for the heat-shock response in intertidal mussels (genus Mytilus) changes as a function of thermal history. Journal of Experimental Biology, 204(20), 3571–3579. 10.1242/jeb.204.20.3571

Carrington, E., Moeser, G. M., Thompson, S. B., Coutts, L. C., & Craig, C. A. (2008). Mussel attachment on rocky shores: The effect of flow on byssus production. Integrative and Comparative Biology, 48(6), 801–807. 10.1093/icb/icn078

Cerullo, A. R., Lai, T. Y., Allam, B., Baer, A., Barnes, W. J. P., Barrientos, Z., Deheyn, D. D., Fudge, D. S., Gould, J., Harrington, M. J., Holford, M., Hung, C.-S., Jain, G., Mayer, G., Medina, M., Monge-Nájera, J., Napolitano, T., Espinosa, E. P., Schmidt, S., … Braunschweig, A. B. (2020). Comparative Animal Mucomics: Inspiration for Functional Materials from Ubiquitous and Understudied Biopolymers. ACS Biomaterials Science & Engineering, 6(10), 5377–5398. 10.1021/acsbiomaterials.0c00713

Chang, M., Ge, J., Liao, M., Rong, X., Wang, Y., Li, B., Li, X., Wang, J., Zhang, Z., Yu, Y., & Wang, C. (2023). Genome-wide DNA methylation and transcription analysis reveal the potential epigenetic mechanism of heat stress response in the sea cucumber Apostichopus japonicus. Frontiers in Marine Science, 10. 10.3389/fmars.2023.1136926

Chapelle, V., & Silvestre, F. (2022). Population Epigenetics: The Extent of DNA Methylation Variation in Wild Animal Populations. Epigenomes, 6(4), 31. 10.3390/epigenomes6040031

Châtel, A., Hamer, B., Jakšić, Ž., Vucelić, V., Talarmin, H., Dorange, G., Schröder, H. C., & Müller, W. E. G. (2011). Induction of apoptosis in mussel Mytilus galloprovincialis gills by model cytotoxic agents. Ecotoxicology, 20(8), 2030–2041. 10.1007/s10646-011-0746-6

Chen, N., Huang, Z., Lu, C., Shen, Y., Luo, X., Ke, C., & You, W. (2019). Different Transcriptomic Responses to Thermal Stress in Heat-Tolerant and Heat-Sensitive Pacific Abalones Indicated by Cardiac Performance. Frontiers in Physiology, 9. 10.3389/fphys.2018.01895

Chen, Y., Pal, B., Visvader, J. E., & Smyth, G. K. (2018). *Differential methylation analysis of reduced representation bisulfite sequencing experiments using edgeR* (No. 6:2055). F1000Research. 10.12688/f1000research.13196.2

Clark, M. S., Thorne, M. A. S., King, M., Hipperson, H., Hoffman, J. I., & Peck, L. S. (2018). Life in the intertidal: Cellular responses, methylation and epigenetics. Functional Ecology, 32(8), 1982–1994. 10.1111/1365-2435.13077

Collins, M., Clark, M. S., & Truebano, M. (2023). The environmental cellular stress response: The intertidal as a multistressor model. Cell Stress and Chaperones, 28(5), 467–475. 10.1007/s12192-023-01348-7

Corfield, A. P. (2015). Mucins: A biologically relevant glycan barrier in mucosal protection. Biochimica Et Biophysica Acta, 1850(1), 236–252. 10.1016/j.bbagen.2014.05.003

Dahlhoff, E. P., Stillman, J. H., & Menge, B. A. (2002). Physiological Community Ecology: Variation in Metabolic Activity of Ecologically Important Rocky Intertidal Invertebrates Along Environmental Gradients1. Integrative and Comparative Biology, 42(4), 862–871. 10.1093/icb/42.4.862

Dainat, J. (2020). *AGAT: Another Gff Analysis Toolkit to handle annotations in any GTF/GFF format* (Version v0.7.0) [Computer software]. https://www.doi.org/10.5281/zenodo.3552717

Deans, C., & Maggert, K. A. (2015). What Do You Mean, “Epigenetic”? Genetics, 199(4), 887–896. 10.1534/genetics.114.173492

Denny, M. W., Dowd, W. W., Bilir, L., & Mach, K. J. (2011). Spreading the risk: Small-scale body temperature variation among intertidal organisms and its implications for species persistence. Journal of Experimental Marine Biology and Ecology, 400(1), 175–190. 10.1016/j.jembe.2011.02.006

Dixon, G., & Matz, M. (2022). Changes in gene body methylation do not correlate with changes in gene expression in Anthozoa or Hexapoda. BMC Genomics, 23(1), 234. 10.1186/s12864-022-08474-z

Dowd, W. W., & Denny, M. W. (In press). Intertidal microcosms of wave-swept rocky shores: Ecological and physiological insights from a uniquely stressful environment. Philosophical Transactions of the Royal Society B.

Dowd, W. W., Felton, C. A., Heymann, H. M., Kost, L. E., & Somero, G. N. (2013). Food availability, more than body temperature, drives correlated shifts in ATP-generating and antioxidant enzyme capacities in a population of intertidal mussels (*Mytilus californianus*). Journal of Experimental Marine Biology and Ecology, 449, 171–185. 10.1016/j.jembe.2013.09.020

Downey-Wall, A. M., Cameron, L. P., Ford, B. M., McNally, E. M., Venkataraman, Y. R., Roberts, S. B., Ries, J. B., & Lotterhos, K. E. (2020). Ocean Acidification Induces Subtle Shifts in Gene Expression and DNA Methylation in Mantle Tissue of the Eastern Oyster (Crassostrea virginica). Frontiers in Marine Science, 7. 10.3389/fmars.2020.566419

Duncan, E. J., Cunningham, C. B., & Dearden, P. K. (2022). Phenotypic Plasticity: What Has DNA Methylation Got to Do with It? Insects, 13(2), 110. 10.3390/insects13020110

Duncan, E. J., Gluckman, P. D., & Dearden, P. K. (2014). Epigenetics, plasticity, and evolution: How do we link epigenetic change to phenotype? Journal of Experimental Zoology Part B: Molecular and Developmental Evolution, 322(4), 208–220. 10.1002/jez.b.22571

Eirin-Lopez, J. M., & Putnam, H. M. (2019). Marine Environmental Epigenetics. In Annual Review of Marine Science (Vol. 11, Issue Volume 11, 2019, pp. 335–368). Annual Reviews. 10.1146/annurev-marine-010318-095114

Ericson, J. A., Laroche, O., Biessy, L., Delorme, N. J., Pochon, X., Thomson-Laing, J., Ragg, N. L. C., & Smith, K. F. (2024). Differential responses of selectively bred mussels (Perna canaliculus) to heat stress—Survival, immunology, gene expression and microbiome diversity. Frontiers in Physiology, 14, 1265879. 10.3389/fphys.2023.1265879

Escobar-Sierra, C., Cañedo-Argüelles, M., Vinyoles, D., & Lampert, K. P. (2024). Unraveling the molecular mechanisms of fish physiological response to freshwater salinization: A comparative multi-tissue transcriptomic study in a river polluted by potash mining. Environmental Pollution, 357, 124400. 10.1016/j.envpol.2024.124400

Ewels, P., Magnusson, M., Lundin, S., & Käller, M. (2016). MultiQC: Summarize analysis results for multiple tools and samples in a single report. Bioinformatics, 32(19), 3047–3048. 10.1093/bioinformatics/btw354

Fitzgerald-Dehoog, L., Browning, J., & Allen, B. J. (2012). Food and Heat Stress in the California Mussel: Evidence for an Energetic Trade-off Between Survival and Growth. The Biological Bulletin, 223(2), 205–216. 10.1086/BBLv223n2p205

Flores, K., Wolschin, F., Corneveaux, J. J., Allen, A. N., Huentelman, M. J., & Amdam, G. V. (2012). Genome-wide association between DNA methylation and alternative splicing in an invertebrate. BMC Genomics, 13(1), 480. 10.1186/1471-2164-13-480

Geeven, G., Zhu, Y., Kim, B. J., Bartholdy, B. A., Yang, S.-M., Macfarlan, T. S., Gifford, W. D., Pfaff, S. L., Verstegen, M. J. A. M., Pinto, H., Vermunt, M. W., Creyghton, M. P., Wijchers, P. J., Stamatoyannopoulos, J. A., Skoultchi, A. I., & de Laat, W. (2015). Local compartment changes and regulatory landscape alterations in histone H1-depleted cells. Genome Biology, 16(1), 289. 10.1186/s13059-015-0857-0

Geyer, K. K., Niazi, U. H., Duval, D., Cosseau, C., Tomlinson, C., Chalmers, I. W., Swain, M. T., Cutress, D. J., Bickham-Wright, U., Munshi, S. E., Grunau, C., Yoshino, T. P., & Hoffmann, K. F. (2017). The Biomphalaria glabrata DNA methylation machinery displays spatial tissue expression, is differentially active in distinct snail populations and is modulated by interactions with Schistosoma mansoni. PLOS Neglected Tropical Diseases, 11(5), e0005246. 10.1371/journal.pntd.0005246

Gleason, L. U., Fekete, F. J., Tanner, R. L., & Dowd, W. W. (2023). Multi-omics reveals largely distinct transcript- and protein-level responses to the environment in an intertidal mussel. The Journal of Experimental Biology, 226(22), jeb245962. 10.1242/jeb.245962

Gleason, L. U., Miller, L. P., Winnikoff, J. R., Somero, G. N., Yancey, P. H., Bratz, D., & Dowd, W. W. (2017). Thermal history and gape of individual Mytilus californianus correlate with oxidative damage and thermoprotective osmolytes. Journal of Experimental Biology, 220(22), 4292–4304. 10.1242/jeb.168450

Gleason, L. U., Strand, E. L., Hizon, B. J., & Dowd, W. W. (2018). Plasticity of thermal tolerance and its relationship with growth rate in juvenile mussels (Mytilus californianus). Proceedings of the Royal Society B: Biological Sciences. 10.1098/rspb.2017.2617

González-Riopedre, M., Novás, A., Dobaño, E., Ramos-Martínez, J. I., & Barcia, R. (2007). Effect of thermal stress on protein expression in the mussel *Mytilus galloprovincialis* Lmk. Comparative Biochemistry and Physiology Part B: Biochemistry and Molecular Biology, 147(3), 531–540. 10.1016/j.cbpb.2007.03.006

Gosling, E. (2015). Marine Bivalve Molluscs. John Wiley & Sons.

Gu, H., Smith, Z. D., Bock, C., Boyle, P., Gnirke, A., & Meissner, A. (2011). Preparation of reduced representation bisulfite sequencing libraries for genome-scale DNA methylation profiling. Nature Protocols, 6(4), 468–481. 10.1038/nprot.2010.190

Guerrero, L., & Bay, R. (2024). Patterns of methylation and transcriptional plasticity during thermal acclimation in a reef-building coral. Evolutionary Applications, 17(7), e13757. 10.1111/eva.13757

Guinle, C., Gurning, R. W., Baratange, C., Cognie, B., Mossion, A., Wielgosz-Collin, G., Bertrand, S., Montiel, G., Poirier, L., Déléris, P., & Zalouk-Vergnoux, A. (2025). Integrating multi-level approaches to assess blue mussel (*Mytilus* spp.) responses to short-term temperature and salinity changes. Marine Environmental Research, 211, 107436. 10.1016/j.marenvres.2025.107436

Haidar, L., Georgescu, M., Drăghici, G. A., Bănățean-Dunea, I., Nica, D. V., & Șerb, A.-F. (2024). DNA Methylation Machinery in Gastropod Mollusks. Life, 14(4), Article 4. 10.3390/life14040537

Halliwell, B., & Gutteridge, J. M. C. (2015). Free Radicals in Biology and Medicine. Oxford University Press.

Harger, R. (1970). The effect of wave impact on some aspects of the biology of sea mussels. Veliger, 12(4), 401–414.

Harley, C. D. G., & Helmuth, B. S. T. (2003). Local- and regional-scale effects of wave exposure, thermal stress, and absolute versus effective shore level on patterns of intertidal zonation. Limnology and Oceanography, 48(4), 1498–1508. 10.4319/lo.2003.48.4.1498

Helmuth, B., Broitman, B. R., Blanchette, C. A., Gilman, S., Halpin, P., Harley, C. D. G., O’Donnell, M. J., Hofmann, G. E., Menge, B., & Strickland, D. (2006). Mosaic Patterns of Thermal Stress in the Rocky Intertidal Zone: Implications for Climate Change. Ecological Monographs, 76(4), 461–479. 10.1890/0012-9615(2006)076%255B0461:MPOTSI%255D2.0.CO;2

Helmuth, B., Harley, C. D. G., Halpin, P. M., O’Donnell, M., Hofmann, G. E., & Blanchette, C. A. (2002). Climate Change and Latitudinal Patterns of Intertidal Thermal Stress. Science, 298(5595), 1015–1017. 10.1126/science.1076814

Helmuth, B. S. T., & Hofmann, G. E. (2001). Microhabitats, Thermal Heterogeneity, and Patterns of Physiological Stress in the Rocky Intertidal Zone. The Biological Bulletin, 201(3), 374–384. 10.2307/1543615

Hiebert, L. S., Soesbe, A., Hsieh, T.-F., Cui, Q., & Yi, S. V. (2025). Integrated Genomic and Methylome Profiling Reveals Promoter Repression and Age-Linked CpGs in the California Mussel. bioRxiv, 2025.08.27.672726. 10.1101/2025.08.27.672726

Hochachka, P. W., & Somero, G. N. (2002). Biochemical Adaptation: Mechanism and Process in Physiological Evolution. Oxford University Press.

Hofmann, G. E., & Somero, G. N. (1995). Evidence for Protein Damage at Environmental Temperatures: Seasonal Changes in Levels of Ubiquitin Conjugates and Hsp70 in the Intertidal Mussel Mytilus trossulus. Journal of Experimental Biology, 198(7), 1509–1518. 10.1242/jeb.198.7.1509

Hofmann, G. E., & Todgham, A. E. (2010). Living in the now: Physiological mechanisms to tolerate a rapidly changing environment. Annual Review of Physiology, 72, 127–145. 10.1146/annurev-physiol-021909-135900

Hu, J., Smith, S. J., Barry, T. N., Jamniczky, H. A., Rogers, S. M., & Barrett, R. D. H. (2021). Heritability of DNA methylation in threespine stickleback (Gasterosteus aculeatus). Genetics, 217(1), iyab001. 10.1093/genetics/iyab001

Huang, Y., Wen, H., Zhang, M., Hu, N., Si, Y., Li, S., & He, F. (2018). The DNA methylation status of *MyoD* and *IGF-I* genes are correlated with muscle growth during different developmental stages of Japanese flounder (*Paralichthys olivaceus*). Comparative Biochemistry and Physiology Part B: Biochemistry and Molecular Biology, 219–220, 33–43. 10.1016/j.cbpb.2018.02.005

Husby, A. (2020). On the Use of Blood Samples for Measuring DNA Methylation in Ecological Epigenetic Studies. Integrative and Comparative Biology, 60(6), 1558–1566. 10.1093/icb/icaa123

Huxtable, R. J. (1992). Physiological actions of taurine. Physiological Reviews, 72(1), 101–163. 10.1152/physrev.1992.72.1.101

Jimenez, A. G., Alves, S., Dallmer, J., Njoo, E., Roa, S., & Dowd, W. W. (2016). Acclimation to elevated emersion temperature has no effect on susceptibility to acute, heat-induced lipid peroxidation in an intertidal mussel (Mytilus californianus). Marine Biology, 163(3), 55. 10.1007/s00227-016-2828-8

Jimenez, A. G., Jayawardene, S., Alves, S., Dallmer, J., & Dowd, W. W. (2015). Micro-scale environmental variation amplifies physiological variation among individual mussels. Proceedings of the Royal Society B: Biological Sciences, 282(1820), 20152273. 10.1098/rspb.2015.2273

Johnson, K. M., & Kelly, M. W. (2020). Population epigenetic divergence exceeds genetic divergence in the Eastern oyster Crassostrea virginica in the Northern Gulf of Mexico. Evolutionary Applications, 13(5), 945–959. 10.1111/eva.12912

Johnson, K. M., Sirovy, K. A., & Kelly, M. W. (2022). Differential DNA methylation across environments has no effect on gene expression in the eastern oyster. Journal of Animal Ecology, 91(6), 1135–1147. 10.1111/1365-2656.13645

Johnson, L. C., Galliart, M. B., Alsdurf, J. D., Maricle, B. R., Baer, S. G., Bello, N. M., Gibson, D. J., & Smith, A. B. (2022). Reciprocal transplant gardens as gold standard to detect local adaptation in grassland species: New opportunities moving into the 21st century. Journal of Ecology, 110(5), 1054–1071. 10.1111/1365-2745.13695

Kauffmann, A., Gentleman, R., & Huber, W. (2009). arrayQualityMetrics—A bioconductor package for quality assessment of microarray data. Bioinformatics, 25(3), 415–416. 10.1093/bioinformatics/btn647

Keller, T. E., Han, P., & Yi, S. V. (2016). Evolutionary Transition of Promoter and Gene Body DNA Methylation across Invertebrate–Vertebrate Boundary. Molecular Biology and Evolution, 33(4), 1019–1028. 10.1093/molbev/msv345

Kho, J., & Ruzzante, D. E. (2024). The role of DNA methylation in facilitating life history trait diversity in fishes. Reviews in Fish Biology and Fisheries, 34(4), 1531–1566. 10.1007/s11160-024-09887-7

Klughammer, J., Romanovskaia, D., Nemc, A., Posautz, A., Seid, C. A., Schuster, L. C., Keinath, M. C., Lugo Ramos, J. S., Kosack, L., Evankow, A., Printz, D., Kirchberger, S., Ergüner, B., Datlinger, P., Fortelny, N., Schmidl, C., Farlik, M., Skjærven, K., Bergthaler, A., … Bock, C. (2023). Comparative analysis of genome-scale, base-resolution DNA methylation profiles across 580 animal species. Nature Communications, 14(1), 232. 10.1038/s41467-022-34828-y

Krueger, F., & Andrews, S. R. (2011). Bismark: A flexible aligner and methylation caller for Bisulfite-Seq applications. Bioinformatics, 27(11), 1571–1572. 10.1093/bioinformatics/btr167

Kültz, D. (2003). Evolution of the cellular stress proteome: From monophyletic origin to ubiquitous function. The Journal of Experimental Biology, 206(Pt 18), 3119–3124. 10.1242/jeb.00549

Kültz, D. (2005). Molecular and evolutionary basis of the cellular stress response. Annual Review of Physiology, 67, 225–257. 10.1146/annurev.physiol.67.040403.103635

Lang, R. P., Bayne, C. J., Camara, M. D., Cunningham, C., Jenny, M. J., & Langdon, C. J. (2009). Transcriptome Profiling of Selectively Bred Pacific Oyster Crassostrea gigas Families that Differ in Tolerance of Heat Shock. Marine Biotechnology (New York, N.Y.), 11(5), 650–668. 10.1007/s10126-009-9181-6

Lawrence, M., Huber, W., Pagès, H., Aboyoun, P., Carlson, M., Gentleman, R., Morgan, M. T., & Carey, V. J. (2013). Software for Computing and Annotating Genomic Ranges. PLOS Computational Biology, 9(8), e1003118. 10.1371/journal.pcbi.1003118

Leeuwis, R. H. J., & Gamperl, A. K. (2022). Adaptations and Plastic Phenotypic Responses of Marine Animals to the Environmental Challenges of the High Intertidal Zone. In Oceanography and Marine Biology. CRC Press.

Levinton, J. S., & Suchanek, T. H. (1978). Geographic variation, niche breadth and genetic differentiation at different geographic scales in the mussels Mytilus californianus and M. edulis. Marine Biology, 49(4), 363–375. 10.1007/BF00455031

Li, J., Zhang, Y., Mao, F., Tong, Y., Liu, Y., Zhang, Y., & Yu, Z. (2017). Characterization and Identification of Differentially Expressed Genes Involved in Thermal Adaptation of the Hong Kong Oyster Crassostrea hongkongensis by Digital Gene Expression Profiling. Frontiers in Marine Science, 4. 10.3389/fmars.2017.00112

Li, Y., Guan, Y., Li, Q., & He, M. (2015). Analysis of DNA methylation in tissues and development stages of pearl oyster Pinctada fucata. Genes & Genomics, 37(3), 263–270. 10.1007/s13258-014-0246-1

Li, Y., Liew, Y. J., Cui, G., Cziesielski, M. J., Zahran, N., Michell, C. T., Voolstra, C. R., & Aranda, M. (2018). DNA methylation regulates transcriptional homeostasis of algal endosymbiosis in the coral model Aiptasia. Science Advances, 4(8), eaat2142. 10.1126/sciadv.aat2142

Li, Y.-F., Yang, X.-Y., Cheng, Z.-Y., Wang, L.-Y., Wang, W.-X., Liang, X., & Yang, J.-L. (2020). Near-future levels of ocean temperature weaken the byssus production and performance of the mussel Mytilus coruscus. Science of The Total Environment, 733, 139347. 10.1016/j.scitotenv.2020.139347

Liew, Y. J., Zoccola, D., Li, Y., Tambutté, E., Venn, A. A., Michell, C. T., Cui, G., Deutekom, E. S., Kaandorp, J. A., Voolstra, C. R., Forêt, S., Allemand, D., Tambutté, S., & Aranda, M. (2018). Epigenome-associated phenotypic acclimatization to ocean acidification in a reef-building coral. Science Advances, 4(6), eaar8028. 10.1126/sciadv.aar8028

Lim, Y.-K., Cheung, K., Dang, X., Roberts, S. B., Wang, X., & Thiyagarajan, V. (2021). DNA methylation changes in response to ocean acidification at the time of larval metamorphosis in the edible oyster, Crassostrea hongkongensis. Marine Environmental Research, 163, 105217. 10.1016/j.marenvres.2020.105217

Liu, A., Zeng, F., Wang, L., Zhen, H., Xia, X., Pei, H., Dong, C., Zhang, Y., & Ding, J. (2023). High temperature influences DNA methylation and transcriptional profiles in sea urchins (Strongylocentrotus intermedius). BMC Genomics, 24(1), 491. 10.1186/s12864-023-09616-7

Lockwood, B. L., & Somero, G. N. (2011). Transcriptomic responses to salinity stress in invasive and native blue mussels (genus Mytilus). Molecular Ecology, 20(3), 517–529. 10.1111/j.1365-294X.2010.04973.x

Lopez-Duarte, P. C., Carson, H. S., Cook, G. S., Fodrie, F. J., Becker, B. J., DiBacco, C., & Levin, L. A. (2012). What Controls Connectivity? An Empirical, Multi-Species Approach. Integrative and Comparative Biology, 52(4), 511–524. 10.1093/icb/ics104

McQuaid, C. D., & Branch, G. M. (1985). Trophic structure of rocky intertidal communities: Response to wave action and implications for energy flow. Marine Ecology Progress Series, 22(2), 153–161.

Moore, L. D., Le, T., & Fan, G. (2013). DNA Methylation and Its Basic Function. Neuropsychopharmacology, 38(1), 23–38. 10.1038/npp.2012.112

Ng, H.-H., & Adrian, B. (1999). DNA methylation and chromatin modification. Current Opinion in Genetics & Development, 9(2), 158–163. 10.1016/S0959-437X(99)80024-0

Nguyen, T. P. L., Nguyen, V. T. A., Do, T. T. T., Nguyen Quang, T., Pham, Q. L., & Le, T. T. (2020). Fatty Acid Composition, Phospholipid Molecules, and Bioactivities of Lipids of the Mud Crab Scylla paramamosain. Journal of Chemistry, 2020(1), 8651453. 10.1155/2020/8651453

Oksanen, J., Simpson, G., Blanchet, F., Kindt, R., Legendre, P., Minchin, P., O’Hara, R., Solymos, P., Stevens, M., Szoecs, E., Wagner, H., Barbour, M., Bedward, M., Bolker, B., Borcard, D., Carvalho, G., Chirico, M., De Caceres, M., Durand, S., … Borman, T. (2025). *vegan: Community Ecology Package* (Version R package version 2.6-10) [Computer software]. https://CRAN.R-project.org/package=vegan

Oldroyd, B. P., & Yagound, B. (2021). The role of epigenetics, particularly DNA methylation, in the evolution of caste in insect societies. Philosophical Transactions of the Royal Society B: Biological Sciences, 376(1826), 20200115. 10.1098/rstb.2020.0115

Ort, B. S., & Pogson, G. H. (2007). Molecular Population Genetics of the Male and Female Mitochondrial DNA Molecules of the California Sea Mussel, Mytilus californianus. Genetics, 177(2), 1087–1099. 10.1534/genetics.107.072934

Padilla, D. K., & Savedo, M. M. (2013). A Systematic Review of Phenotypic Plasticity in Marine Invertebrate and Plant Systems. In Advances in Marine Biology (Vol. 65, pp. 67–94). Elsevier. 10.1016/B978-0-12-410498-3.00002-1

Paggeot, L. X., DeBiasse, M. B., Escalona, M., Fairbairn, C., Marimuthu, M. P. A., Nguyen, O., Sahasrabudhe, R., & Dawson, M. N. (2022). Reference genome for the California ribbed mussel, *Mytilus californianus*, an ecosystem engineer. Journal of Heredity, 113(6), 681–688. 10.1093/jhered/esac041

Paine, R. T. (1974). Intertidal community structure. Oecologia, 15(2), 93–120. 10.1007/BF00345739

Pales Espinosa, E., Koller, A., & Allam, B. (2016). Proteomic characterization of mucosal secretions in the eastern oyster, *Crassostrea virginica*. Journal of Proteomics, 132, 63–76. 10.1016/j.jprot.2015.11.018

Pan, Y., Chen, L., Cheng, J., Zhu, X., Wu, P., Bao, L., Chu, W., He, S., Liang, X., & Zhang, J. (2021). Genome-wide DNA methylation profiles provide insight into epigenetic regulation of red and white muscle development in Chinese perch *Siniperca chuatsi*. Comparative Biochemistry and Physiology Part B: Biochemistry and Molecular Biology, 256, 110647. 10.1016/j.cbpb.2021.110647

Park, K., Lee, J. S., Kang, J.-C., Kim, J. W., & Kwak, I.-S. (2015). Cascading effects from survival to physiological activities, and gene expression of heat shock protein 90 on the abalone Haliotis discus hannai responding to continuous thermal stress. Fish & Shellfish Immunology, 42(2), 233–240. 10.1016/j.fsi.2014.10.036

Pereira, T. M., Minari, M., Carvajalino-Fernández, J. M., Moreira, D. C., & Hermes-Lima, M. (2025). Redox Metabolism During Aerial Exposure of the Sea Urchin Echinometra lucunter: An Ecophysiological Perspective. Animals, 15(9), 1251. 10.3390/ani15091251

Pfister, C. A., Gilbert, J. A., & Gibbons, S. M. (2014). The role of macrobiota in structuring microbial communities along rocky shores. PeerJ, 2, e631. 10.7717/peerj.631

R Core Team. (2023). R: A Language and Environment for Statistical Computing [Computer software]. R Foundation for Statistical Computing, Vienna, Austria. https://www.R-project.org/

Roberto, A.-E., Ana M., I., Steven, R. B., Maria Teresa, S.-G., & Cristina, E.-F. (2021). Differentially methylated gene regions between resistant and susceptible heat-phenotypes of the Pacific oyster Crassostrea gigas. Aquaculture, 543, 736923. 10.1016/j.aquaculture.2021.736923

Robinson, M. D., McCarthy, D. J., & Smyth, G. K. (2010). edgeR: A Bioconductor package for differential expression analysis of digital gene expression data. Bioinformatics, 26(1), 139–140. 10.1093/bioinformatics/btp616

Rossi, F., Palombella, S., Pirrone, C., Mancini, G., Bernardini, G., & Gornati, R. (2016). Evaluation of tissue morphology and gene expression as biomarkers of pollution in mussel *Mytilus galloprovincialis* caging experiment. Aquatic Toxicology, 181, 57–66. 10.1016/j.aquatox.2016.10.018

Sagarin, R. D., Barry, J. P., Gilman, S. E., & Baxter, C. H. (1999). Climate-Related Change in an Intertidal Community over Short and Long Time Scales. Ecological Monographs, 69(4), 465–490. 10.1890/0012-9615(1999)069%255B0465:CRCIAI%255D2.0.CO;2

Seed, R., & Suchanek, T. (1992). Population and communtiy ecology of Mytilus (pp. 87–169).

Seiler Vellame, D., Castanho, I., Dahir, A., Mill, J., & Hannon, E. (2021). Characterizing the properties of bisulfite sequencing data: Maximizing power and sensitivity to identify between-group differences in DNA methylation. BMC Genomics, 22(1), 446. 10.1186/s12864-021-07721-z

Silliman, K., Spencer, L. H., White, S. J., & Roberts, S. B. (2023). Epigenetic and Genetic Population Structure is Coupled in a Marine Invertebrate. Genome Biology and Evolution, 15(2), evad013. 10.1093/gbe/evad013

Somero, G., Lockwood, B. L., & Tomanek, L. (2016). Biochemical Adaptation: Response to Environmental Challenges, from Life’s Origins to the Anthropocene. Palgrave Macmillan.

Somero, G. N. (2010). The physiology of climate change: How potentials for acclimatization and genetic adaptation will determine ‘winners’ and ‘losers.’ Journal of Experimental Biology, 213(6), 912–920. 10.1242/jeb.037473

Sork, V. L. (2018). Genomic Studies of Local Adaptation in Natural Plant Populations. Journal of Heredity, 109(1), 3–15. 10.1093/jhered/esx091

Stillman, J. H., Amri, A. B., Holdreith, J. M., Hooper, A., Leon, R. V., Pruett, L. R., & Bukaty, B. M. (2025). Ecophysiological responses to heat waves in the marine intertidal zone. Journal of Experimental Biology, 228(2), JEB246503. 10.1242/jeb.246503

Stillman, J. H., & Somero, G. N. (2000). A Comparative Analysis of the Upper Thermal Tolerance Limits of Eastern Pacific Porcelain Crabs, Genus *Petrolisthes*: Influences of Latitude, Vertical Zonation, Acclimation, and Phylogeny. Physiological and Biochemical Zoology, 73(2), 200–208. 10.1086/316738

Sun, M., Yang, Z., Liu, L., & Duan, L. (2022). DNA Methylation in Plant Responses and Adaption to Abiotic Stresses. International Journal of Molecular Sciences, 23(13), 6910. 10.3390/ijms23136910

Suzuki, M. M., & Bird, A. (2008). DNA methylation landscapes: Provocative insights from epigenomics. Nature Reviews Genetics, 9(6), 465–476. 10.1038/nrg2341

Tan, C., Shi, C., Li, Y., Teng, W., Li, Y., Fu, H., Ren, L., Yu, H., Li, Q., & Liu, S. (2022). Comparative Methylome Analysis Reveals Epigenetic Signatures Associated with Growth and Shell Color in the Pacific Oyster, Crassostrea gigas. Marine Biotechnology, 24(5), 911–926. 10.1007/s10126-022-10154-8

Thompson, R. J., & Bayne, B. L. (1972). Active metabolism associated with feeding in the mussel Mytilus edulis L. Journal of Experimental Marine Biology and Ecology, 9(1), 111–124. 10.1016/0022-0981(72)90011-1

Tomanek, L., & Zuzow, M. J. (2010). The proteomic response of the mussel congeners Mytilus galloprovincialis and M. trossulus to acute heat stress: Implications for thermal tolerance limits and metabolic costs of thermal stress. The Journal of Experimental Biology, 213(Pt 20), 3559–3574. 10.1242/jeb.041228

Tweedie, S., Charlton, J., Clark, V., & Bird, A. (1997). Methylation of genomes and genes at the invertebrate-vertebrate boundary. Molecular and Cellular Biology, 17(3), 1469–1475.

Valdivieso, A., Morga, B., Degremont, L., Mege, M., Courtay, G., Dorant, Y., Escoubas, J.-M., Gawra, J., de Lorgeril, J., Mitta, G., Cosseau, C., & Vidal-Dupiol, J. (2025). DNA methylation landscapes before and after Pacific Oyster Mortality Syndrome are different within and between resistant and susceptible *Magallana gigas*. Science of The Total Environment, 962, 178385. 10.1016/j.scitotenv.2025.178385

Van Buuren, S., & Groothuis-Oudshoorn, K. (2011). **mice**: Multivariate Imputation by Chained Equations in *R*. Journal of Statistical Software, 45(3). 10.18637/jss.v045.i03

van der Veen, J. N., Kennelly, J. P., Wan, S., Vance, J. E., Vance, D. E., & Jacobs, R. L. (2017). The critical role of phosphatidylcholine and phosphatidylethanolamine metabolism in health and disease. Biochimica et Biophysica Acta (BBA) - Biomembranes, 1859(9, Part B), 1558–1572. 10.1016/j.bbamem.2017.04.006

Venkataraman, Y. R., Downey-Wall, A. M., Ries, J., Westfield, I., White, S. J., Roberts, S. B., & Lotterhos, K. E. (2020). General DNA Methylation Patterns and Environmentally-Induced Differential Methylation in the Eastern Oyster (Crassostrea virginica). Frontiers in Marine Science, 7. 10.3389/fmars.2020.00225

Venkataraman, Y. R., Greiner-Ferris, K., White, S., & Roberts, S. (2024). DNA Methylation Assessment. MarineOmics. https://marineomics.github.io/FUN_02_DNA_methylation.html

Venkataraman, Y. R., Huffmyer, A. S., White, S. J., Downey-Wall, A., Ashey, J., Becker, D. M., Bengtsson, Z., Putnam, H. M., Strand, E., Rodríguez-Casariego, J. A., Wanamaker, S. A., Lotterhos, K. E., & Roberts, S. B. (2024). DNA methylation correlates with transcriptional noise in response to elevated pCO2 in the eastern oyster (*Crassostrea virginica*). Environmental Epigenetics, 10(1), dvae018. 10.1093/eep/dvae018

Venkataraman, Y. R., White, S. J., & Roberts, S. B. (2022). Differential DNA methylation in Pacific oyster reproductive tissue in response to ocean acidification. BMC Genomics, 23(1), 556. 10.1186/s12864-022-08781-5

Vogt, G. (2022). Studying phenotypic variation and DNA methylation across development, ecology and evolution in the clonal marbled crayfish: A paradigm for investigating epigenotype-phenotype relationships in macro-invertebrates. The Science of Nature, 109(1), 16. 10.1007/s00114-021-01782-6

Walker, S. T., Mantle, D., Bythell, J. C., & Thomason, J. C. (2000). Oxidative-stress: Comparison of species specific and tissue specific effects in the marine bivalves *Mytilus edulis* (L.) and *Dosinia lupinus* (L.). Comparative Biochemistry and Physiology Part B: Biochemistry and Molecular Biology, 127(3), 347–355. 10.1016/S0305-0491(00)00266-2

Wang, J., Ren, R., & Yao, C.-L. (2018). Oxidative stress responses of Mytilus galloprovincialis to acute cold and heat during air exposure. Journal of Molluscan Studies, 84(3), 285–292. 10.1093/mollus/eyy027

Wang, J., Yang, B., Zhang, X., Liu, S., Pan, X., Ma, C., Ma, S., Yu, D., & Wu, W. (2023). Chromobox proteins in cancer: Multifaceted functions and strategies for modulation (Review). International Journal of Oncology, 62(3), 36. 10.3892/ijo.2023.5484

Wang, X., Cong, R., Li, A., Wang, W., Zhang, G., & Li, L. (2023). Transgenerational effects of intertidal environment on physiological phenotypes and DNA methylation in Pacific oysters. Science of The Total Environment, 871, 162112. 10.1016/j.scitotenv.2023.162112

Wang, X., Li, A., Wang, W., Que, H., Zhang, G., & Li, L. (2021). DNA methylation mediates differentiation in thermal responses of Pacific oyster (Crassostrea gigas) derived from different tidal levels. Heredity, 126(1), 10–22. 10.1038/s41437-020-0351-7

Watson, H., Videvall, E., Andersson, M. N., & Isaksson, C. (2017). Transcriptome analysis of a wild bird reveals physiological responses to the urban environment. Scientific Reports, 7(1), 44180. 10.1038/srep44180

West-Eberhard, M. J. (1989). Phenotypic Plasticity and the Origins of Diversity. Annual Review of Ecology and Systematics, 20, 249–278.

Williams, E. E., & Somero, G. N. (1996). Seasonal-, Tidal-Cycle- and Microhabitat-Related Variation in Membrane Order of Phospholipid Vesicles from Gills of the Intertidal Mussel Mytilus Californianus. Journal of Experimental Biology, 199(7), 1587–1596. 10.1242/jeb.199.7.1587

Wu, F., Sokolov, E. P., Khomich, A., Fettkenhauer, C., Schnell, G., Seitz, H., & Sokolova, I. M. (2022). Interactive effects of ZnO nanoparticles and temperature on molecular and cellular stress responses of the blue mussel *Mytilus edulis*. Science of The Total Environment, 818, 151785. 10.1016/j.scitotenv.2021.151785

Young, M. D., Wakefield, M. J., Smyth, G. K., & Oshlack, A. (2010). Gene ontology analysis for RNA-seq: Accounting for selection bias. Genome Biology, 11(2), R14. 10.1186/gb-2010-11-2-r14

Zhang, H., Tan, K., Li, S., Ma, H., & Zheng, H. (2020). DNA methylation in molluscs growth and development: An overview. Aquaculture Research, 53(14), 4893–4900. 10.1111/are.14966

Zhang, M., Zhao, J., Li, A., Zhao, M., Huo, M., Deng, J., Wang, L., Wang, W., Zhang, G., & Li, L. (2025). LncRNA-Mediated Tissue-Specific Plastic Responses to Salinity Changes in Oysters. International Journal of Molecular Sciences, 26(10), 4523. 10.3390/ijms26104523

Zhang, W., Storey, K. B., & Dong, Y. (2021). Synchronization of seasonal acclimatization and short-term heat hardening improves physiological resilience in a changing climate. Functional Ecology, 35(3), 686–695. 10.1111/1365-2435.13768

Zhang, X., Li, Q., Kong, L., & Yu, H. (2017). DNA methylation changes detected by methylation-sensitive amplified polymorphism in the Pacific oyster (Crassostrea gigas) in response to salinity stress. Genes & Genomics, 39(11), 1173–1181. 10.1007/s13258-017-0583-y

Zhou, Y., Xu, L., Bickhart, D. M., abdel Hay, E. H., Schroeder, S. G., Connor, E. E., Alexander, L. J., Sonstegard, T. S., Van Tassell, C. P., Chen, H., & Liu, G. E. (2016). Reduced representation bisulphite sequencing of ten bovine somatic tissues reveals DNA methylation patterns and their impacts on gene expression. BMC Genomics, 17, 779. 10.1186/s12864-016-3116-1

