## Supplemental Materials for "Tissue-specific plasticity of DNA methylation across intertidal microhabitats in juvenile mussels *(Mytilus californianus)*"

**Affiliations:**
[1]Department of Ecology and Evolutionary Biology, University of California, Santa Cruz, Santa Cruz, CA
[2]Environmental Science & Policy Program, Chapman University, Orange, CA
[3]School of Biological Sciences, Washington State University, Pullman, WA

**Table of Contents**

Supplemental Text -----------------------------------------------------------------------1
References ----------------------------------------------------------------------- 5
Supplemental Table Legends -------------------------------------------------------------6
Supplemental Figures ------------------------------------------------------------------ 8

### Supplemental Text

#### Outlier sample removal and analysis of percent methylation in a CpG context

Outlier samples were identified separately among the foot (33 samples) and gill (32 samples) tissue subsets. Data was imputed for CpG sites with missing data. Only CpG sites with a minimum coverage of three reads in at least 22 individuals were retained in the matrix for imputation. For each tissue type, the imputed methylation proportion matrix was used as input for *arrayQualityMetrics*. We screened for significant outlier samples using the *arrayQualityMetrics* distance-between-arrays test, which detects outliers in PCA space (Kauffmann et al., 2009). We removed three significant outlier gill samples (15W-G_S12, 60G-G_S58, 38B-G_S78) for all subsequent analyses; no outliers were detected in the foot samples.

To assess whether levels of DNA methylation across all CpG sites changed among treatments and between tissues, the proportion of methylation calls from all read alignments to CpGs was extracted from the Bismark summary reports (the '% methylation in a CpG context' metric) for the 40 retained samples after filtering (Krueger & Andrews, 2011).This value is distinct from the genome-wide mean CpG methylation described in the main text, as it is the proportion of methylation calls from the alignment rather than a mean proportion of methylation across CpGs. This alignment-based % CpG methylation metric helped assess whether samples exhibited differences in coverage of methylated or unmethylated regions, or if samples potentially differed in bisulfite conversion of reads.

#### Model comparison in differential methylation (DM) analyses

DM analyses were performed with the *R* package *edgeR* v4.0.16 (Robinson et al., 2010) using coverage files generated from Bismark. We fitted separate models to gill and foot data. We first used the *edgeR glmFit* function to fit negative binomial generalized linear models that predicted variation in methylated and unmethylated read counts as a function of fixed effects for origin site, transplant site, and the shell length of the mussels at time of sampling. DM CpGs associated with transplant site, origin site, and shell length were identified by conducting likelihood ratio tests with the *glmLRT* function. The *edgeR* tagwise dispersion function was used to calculate CpG-specific coefficients of variation for negative binomial models of DM.

We additionally fitted a model of differential methylation with the same fixed effects as described in the main text (origin site, transplant site, and shell length) and included an interaction term between the origin and transplant site. However, log2FC values that were generated in response to the transplant effect were extremely and unrealistically high (log2FC > 50 in many cases), an issue likely caused by model overfitting: small sample sizes in origin x transplant site groups likely permitted spuriously large log2FC values. Adding an intercept to the negative binomial GLMs of DM mitigated inflated log2FC DM coefficients. To evaluate the influence of the additional interaction term on the original model, we therefore compared our non-intercept model with only main effects (reported in the main text) to an alternative model with an intercept and an origin x transplant interaction term. We determined whether the interaction term explained a large amount of DM unobserved by the non-intercept model by screening for DM using methods described in the main text.

To determine whether the interaction term altered coefficients for DM associated with origin and transplant sites, we measured the correlation in log2FC DM coefficients predicted by the non-intercept and intercept model using Pearson correlation. We found that the log2FC values generated from the two models were highly correlated (mean correlation: 0.72; $p < 0.001$; Figure S4), and the addition of the interaction term had minimal effects on model predictions. We also assessed the number of DM CpGs detected associated with the interaction term from the interaction model and only found 27 DM CpGs in foot (8 hypomethylated and 19 hypermethylated) and 11 DM CpGs in gill (5 hypomethylated and 6 hypermethylated), respectively (Figure S5). Given the low number of DM CpG sites associated with the interaction term and the relatively high correlation between the two models, the interaction model was not used in our final analyses to prevent the risk of overfitting. While we acknowledge that the effects of environmental exposure on DNA methylation can be dependent on the origins of the mussels, significantly larger sample sizes would increase confidence in the outputs of a more complex statistical model.

Significant differences in the prevalence of CpG DM between genomic features (e.g. exons, introns, promoters, untranslated regions (UTRs), and intergenic regions) were evaluated using

GLMs with a binomial distribution and logit link. GLMs were separately fit for DM associated with origin, transplant site, and in each tissue (four total models). The resulting p-value from each pair-wise comparison was adjusted for multiple testing using the Tukey method with the R package *emmeans* (Lenth, 2025).

**GO enrichment analyses**

Gene Ontology (GO) enrichment analysis was performed separately for each tissue type to investigate the functions of the DM genes using the *R* package *GOseq* v1.54.0 (Young et al., 2010) and a rank-based Gene Ontology Analysis with Adaptive Clustering and Mann-Whitney U test based on ranking of mean gene-wide log2-fold-changes of DM (GO-MWU, https://github.com/z0on/GO_MWU). *GOseq* can account for bias introduced by differences in gene length during enrichment analyses. Two inputs were required: a background list of genes that were measured and a binary vector indicating genes that were differentially expressed. GO annotations for *M. californianus* were obtained from the reference genome annotation xbMytCali1.0.p from NCBI (GCF_021869535.1; 69,518 GO terms). Due to the specificity of the GO annotations for each gene, we extracted the parent GO terms (one level up in the GO hierarchy) for all the NCBI GO annotations using the *R* packages *GO.db* v3.18.0 and *AnnotationDbi* v1.64.1 (Carlson, 2023; Pagès et al., 2023). This step generated 2,105 parent GO terms for all downstream analyses from the list of 2,032 original GO terms. A background list of genes for GO enrichment analyses was obtained by extracting CpG sites analyzed in the DM analyses that overlap with annotated *M. californianus* genes. We then matched all CpG-overlapping genes with their respective GO terms. GO terms that contained fewer than three genes were not considered. To account for the possibility that the likelihood of a gene being characterized as differentially methylated is influenced by the number of associated CpG sites, a matrix containing the number of CpGs associated with each gene was supplied for bias correction. Overrepresented biological processes, molecular functions, and cellular components were identified with an FDR-adjusted threshold of alpha-value < 0.05 (Benjamini & Hochberg, 1995). For origin and transplant site effect in foot, 15 and 18 DM genes were tested against a background list of 2,793 genes, representing 884 GO terms. For gill, 33 and 40 DM genes were tested against a background list of 2,729 genes, representing 840 GO terms. For foot, there were 12 over-represented GO terms associated with the origin site effect and 14 associated with the transplant site effect (unadjusted $p < 0.05$). For gill, 15 over-represented GO terms were associated with origin site effect, and 11 were associated with transplant site effect. However, these enrichments were not significant after correction for multiple testing (Table S6).

We also ran a rank-based Gene Ontology Analysis with Adaptive Clustering and Mann-Whitney U test based on ranking of mean gene-wide log2-fold-changes of DM (GO-MWU, https://github.com/z0on/GO_MWU). Using the same sets of filtered CpG sites and their associated genes used in *GOseq*, we averaged the log2FC values per gene associated with origin and transplant site effects, generating separate mean logFC scores for each effect for each tissue type. Only GO categories that contained at least 2 genes were considered, and those that contained more than 1% of the total number of genes were not included in the analysis. GO enrichment was tested for biological processes, molecular functions, and cellular components separately. This method tests whether genes belonging to a certain GO category are significantly clustered near the top or the bottom of the global ranked list of genes, rather than being spread evenly. Redundant GO categories were collapsed under the name of the lower-level (more specific) term using cutTreeHeight 0.25, where a group of categories will be merged if the most dissimilar two of them share >75% of genes included in the smaller of the two. No significant enrichment was detected in both tissue types associated with all treatment groups.

**References:**

Benjamini, Y., & Hochberg, Y. (1995). Controlling the False Discovery Rate: A Practical and Powerful Approach to Multiple Testing. *Journal of the Royal Statistical Society: Series B (Methodological)*, *57*(1), 289–300. https://doi.org/10.1111/j.2517-6161.1995.tb02031.x

Kauffmann, A., Gentleman, R., & Huber, W. (2009). arrayQualityMetrics—A bioconductor package for quality assessment of microarray data. *Bioinformatics*, *25*(3), 415–416. https://doi.org/10.1093/bioinformatics/btn647

Krueger, F., & Andrews, S. R. (2011). Bismark: A flexible aligner and methylation caller for Bisulfite-Seq applications. *Bioinformatics*, *27*(11), 1571–1572. https://doi.org/10.1093/bioinformatics/btr167

Robinson, M. D., McCarthy, D. J., & Smyth, G. K. (2010). edgeR: A Bioconductor package for differential expression analysis of digital gene expression data. *Bioinformatics*, *26*(1), 139–140. https://doi.org/10.1093/bioinformatics/btp616

Young, M. D., Wakefield, M. J., Smyth, G. K., & Oshlack, A. (2010). Gene ontology analysis for RNA-seq: Accounting for selection bias. *Genome Biology*, *11*(2), R14. https://doi.org/10.1186/gb-2010-11-2-r14

### Supplemental Table Legends

**Supplemental Table 1:** Sample information and RRBS sequencing summary by lane. Each row represents sequencing results for an individual sample split across four lanes (L001-L004). Samples highlighted in gray were selected in the differential methylation analyses. (n=65 total; after filtering: n=20 per gill and foot subset)

**Supplemental Table 2:** Bismark alignment and RRBS sequencing statistics for concatenated reads per sample. Samples highlighted in gray were selected in the differential methylation analyses. (n=65 total; after filtering: n=20 per gill and foot subset)

**Supplemental Table 3. Summary of statistical tests**

a. Results of linear mixed effect models assessing the relationship between mussel shell length and PC1 and PC2

b. Results of generalized linear mixed effect models with beta distribution testing whether raw and poster-filtered genome-wide methylation levels differed between tissue type, origin site, and transplant treatments

c. Results of Fisher's exact test on DM results assessing whether different genomic features were overrepresented relative to the genomic background. Tests were conducted separately for foot and gill tissues, as well as between tissues. Each row corresponds to one feature tested against all other features combined

d. Results of binomial generalized linear models with logit links testing whether certain genomic features were more likely to contain DM CpGs. Tests were conducted separately for each effect within each tissue type. Promoter regions were excluded from the foot origin model because no DM CpGs were detected in this category

**Supplemental Table 4. Summary of results from DM analyses in foot samples**

a. DM CpGs identified in foot tissue for origin site effect. CpG loci highlighted in orange were shared between origin site and transplant site effects
b. DM CpGs identified in foot tissue for transplant site effect. CpG loci highlighted in orange were shared between origin site and transplant site effects
c. DM CpGs identified in foot tissue for shell length effect

**Supplemental Table 5. Summary of results from DM analyses in gill samples**

a. DM CpGs identified in gill tissue for origin site effect. CpG loci highlighted in orange were shared between origin site and transplant site effects
b. DM CpGs identified in gill tissue for transplant site effect. CpG loci highlighted in orange were shared between origin site and transplant site effects
c. DM CpGs identified in gill tissue for shell length effect

**Supplemental Table 6. Summary of results from GO enrichment analysis (goseq)**

a. GO enrichment analysis for DM genes in response to origin effect in foot
b. GO enrichment analysis for DM genes in response to transplant effect in foot
c. GO enrichment analysis for DM genes in response to origin effect in gill
d. GO enrichment analysis for DM genes in response to transplant effect in gill

### Supplemental Figures

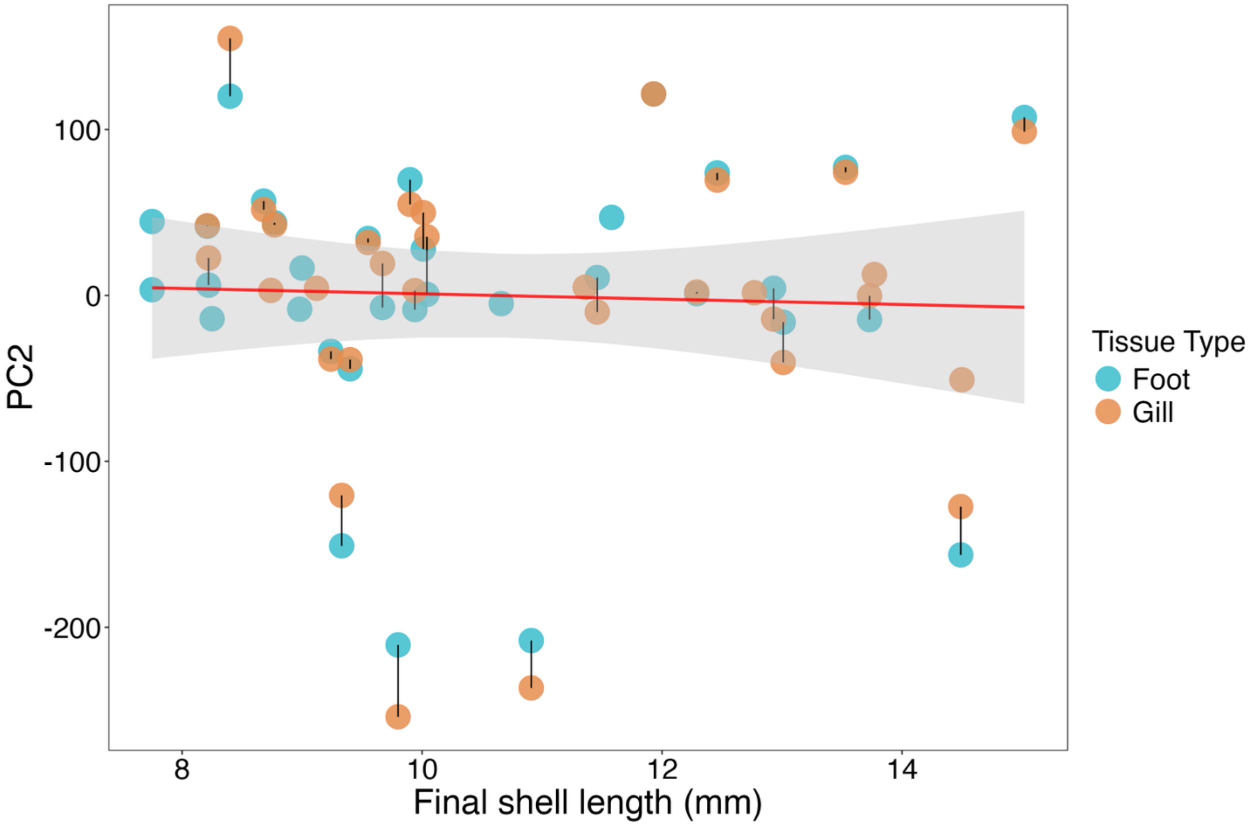


**Figure S1.** Correlation between PC2 of DNA methylation data and shell length of *M. californianus* was not significant (p = 0.79). CpG sites with missing data were imputed. Tissues collected from the same individual are connected with a black line. The red line represents the best-fit line, and ribbons represent the 95% confidence interval. Blue and orange dots represent foot and gill tissue samples, respectively. N = 33 for foot and N = 32 for gill.

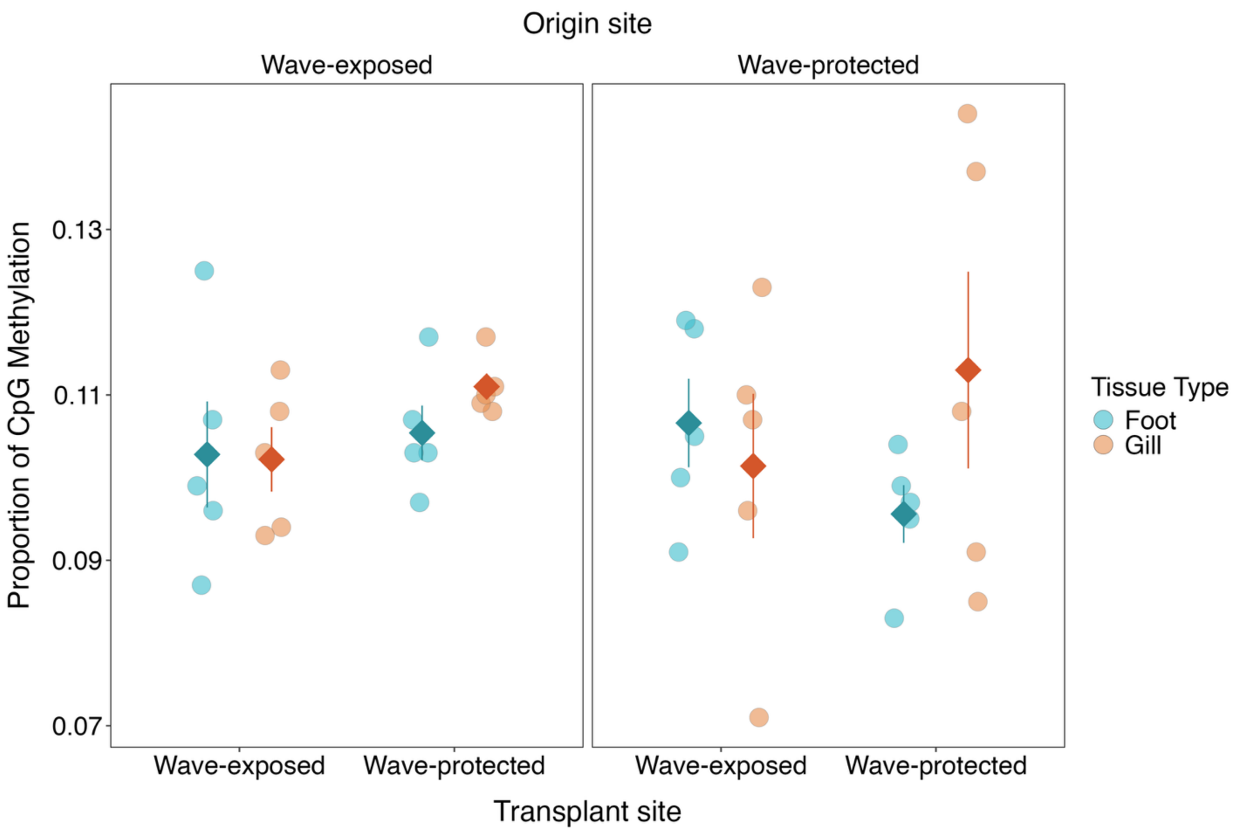


**Figure S2.** Genome-wide DNA methylation levels (the percent methylation of reads aligned to CpGs) did not significantly differ between tissue type, origin site, or transplant site effects. Juvenile *M. californianus* were transplanted between cool, wave-exposed and warm, wave-protected microhabitats. Each point represents a sample. Diamonds represent the mean methylation proportion, and the whiskers represent ± SE.

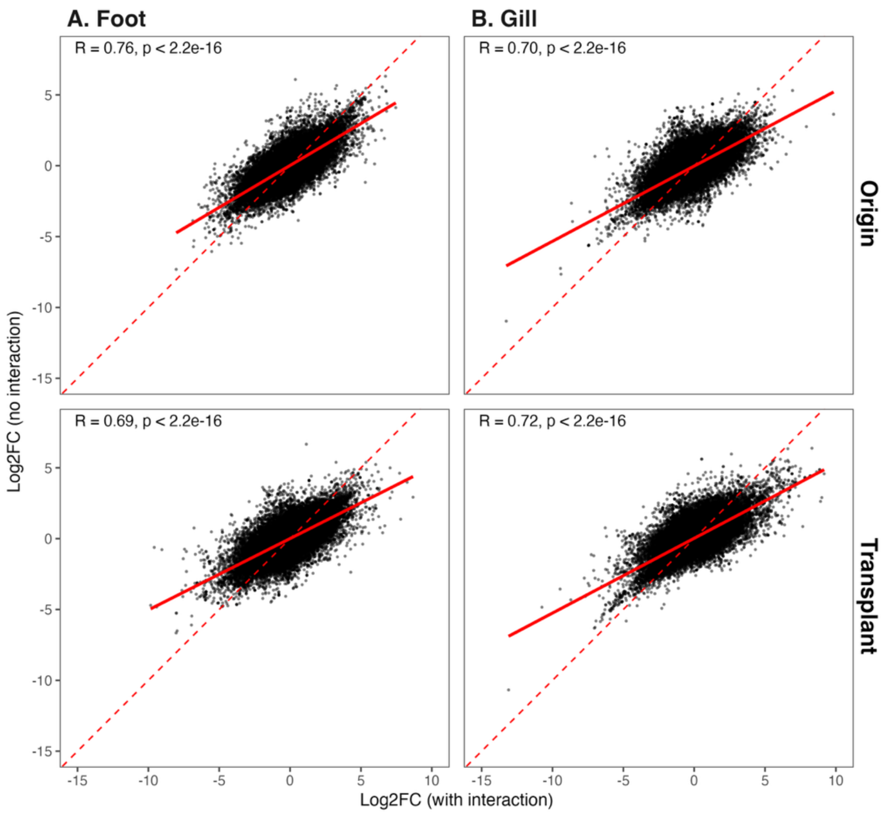


**Figure S3.** Positive and significant correlation in the log2FC values between the edgeR model with and without the interaction term associated with origin and transplant site effects in **A**) foot, and **B)** gill (Pearson's correlation tests, $p < 0.001$). Each point represents a shared CpG site between the two models associated with origin or transplant site effect in foot ($n = 77,434$) or gill ($n = 76,636$). The red solid lines represent the lines of best fit. The red diagonal dashed line represents a 1:1 relationship between Log2FC values with and without an interaction term.

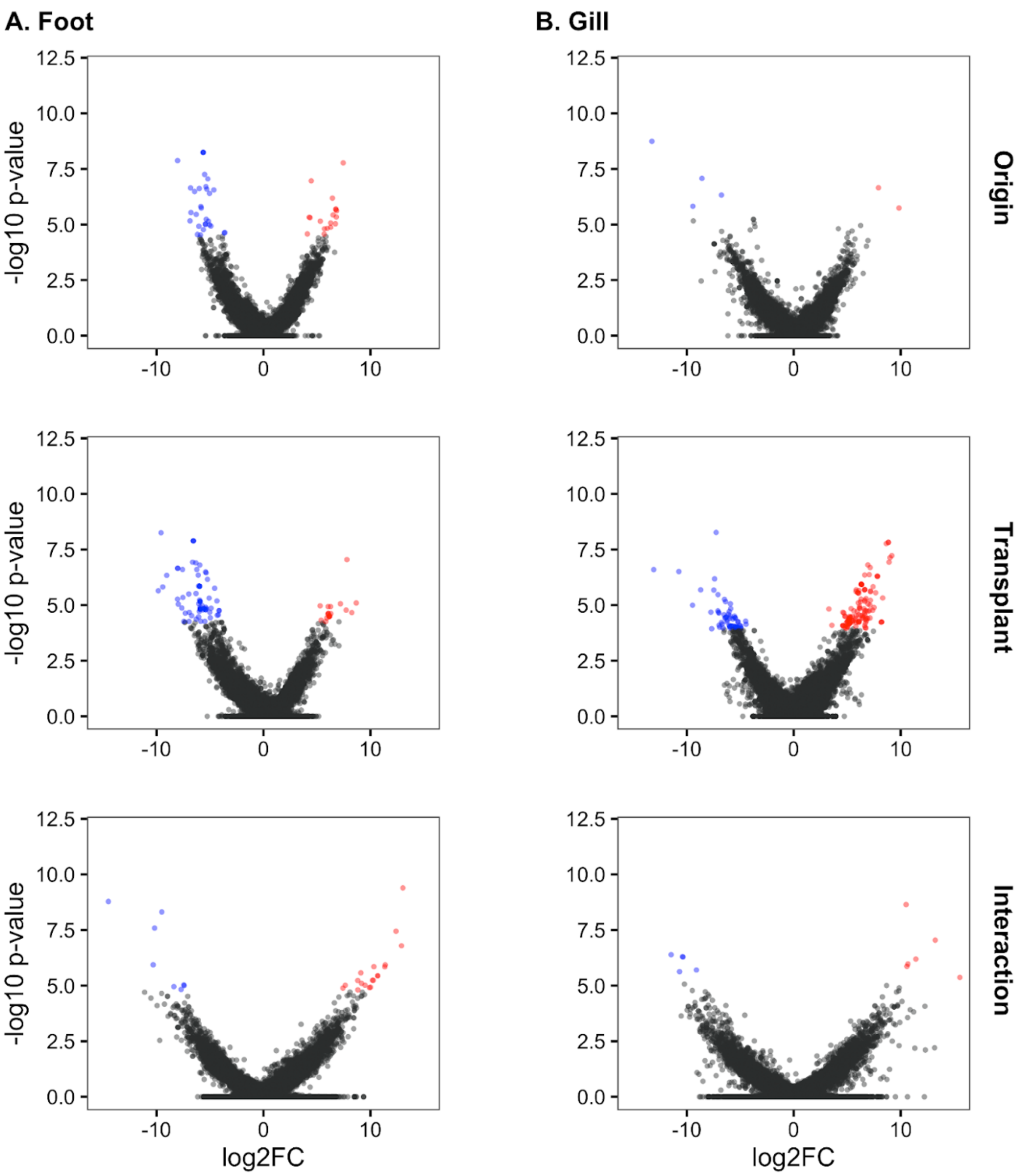


**Figure S4.** Differentially methylated CpGs identified based on the effect of origin site (top) and transplant site (middle), and the interaction between origin and transplant site (bottom) for **A.** foot and **B.** gill samples. Each dot represents a CpG locus. hyper- (red) and hypo (blue)-methylated CpGs, respectively, colored based on an FDR-adjusted p-value of $< 0.05$. Exposed sites were used as a reference for each comparison.

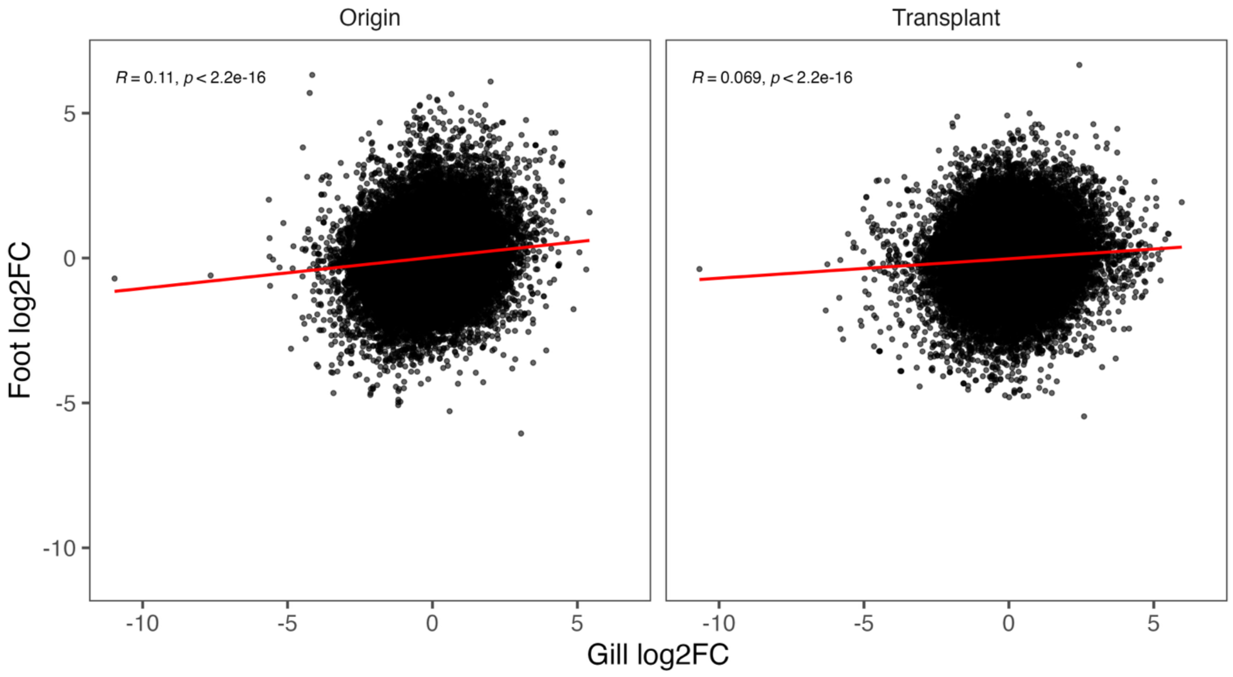


**Figure S5.** Weakly positive and significant correlation in the log2FC values between foot and gill in response to origin and transplant effect (Pearson's correlation, origin r = 0.11, transplant r = 0.069, p < 0.001 for each). Each point represents a shared CpG site between foot and gill (n = 63,586). The red lines represent the lines of best fit.
